# Computational Modeling of the Gut Microbiota Predicts Metabolic Mechanisms of Recurrent *Clostridioides difficile* Infection

**DOI:** 10.1101/2020.04.10.036111

**Authors:** Michael A. Henson

## Abstract

Approximately 30% of patients who have a *Clostridioides difficile* infection (CDI) will suffer at least one incident of reinfection. While the underlying causes of CDI recurrence are poorly understood, interactions between *C. difficile* and other commensal gut bacteria are thought to play an important role. In this study, an *in silico* metagenomics pipeline was used to process taxa abundance data from 225 CDI patient stool samples into sample-specific models of bacterial community metabolism. The predicted metabolite production capabilities of each community were shown to provide improved recurrence prediction compared to direct use of taxa abundance data. More specifically, clustered metabolite synthesis rates generated from post-diagnosis samples produced a high *Enterobacteriaceae* cluster with disproportionate numbers of recurrent samples and patients. This cluster was predicted to have significantly reduced capabilities for secondary bile acid synthesis but elevated capabilities for aromatic amino acid catabolism. When applied to 40 samples from fecal microbiota transplantation (FMT) patients and their donors, community modeling generated a high *Enterobacteriaceae* cluster with a disproportionate number of pre-FMT samples. This cluster also was predicted to exhibit reduced secondary bile acid synthesis and elevated aromatic amino acid catabolism. Because clustering of CDI and FMT samples did not identify statistical differences in *C. difficile* abundances, these model predictions support the hypothesis that *Enterobacteriaceae* may create a gut environment favorable for *C. difficile* spore germination and toxin synthesis.

**Importance:** *Clostridioides difficile* is an opportunistic human pathogen responsible for acute and sometimes chronic infections of the colon. Elderly individuals who are immunocompromised, frequently hospitalized and recipients of antibiotics are particular susceptible to infection. Approximately 30% of treated patients will suffer at least one episode of reinfection, commonly termed recurrence. The objective of the current study was to utilize computational metabolic modeling to investigate the hypothesis that recurrent infections are related to the composition of the gut bacterial community within each patient. Our model predictions suggest that patients who have high compositions of the bacterial family *Enterobacteriaceae* during antibiotic treatment are more likely to develop recurrent infections due to a metabolically-disrupted gut environment. Successful treatment of recurrent patients with transplanted fecal matter is predicted to correct this metabolic disruption, suggesting that interactions between *C. difficile* and *Enterobacteriaceae* are worthy of additional study.

## Introduction

The anaerobic bacterium *Clostridioides difficile* is an opportunistic pathogen responsible for infections, primarily in the human colon (1). *C. difficile* infection (CDI) is most common in elderly patients previously treated with broad spectrum antibiotics that disrupt the healthy gut microbiota and produce a dysbiotic environment conducive to *C. difficile* germination, expansion and pathogenicity (2, 3). CDI has become particularly common in hospital settings due to the ability of *C. difficile* to form spores that adhere to surfaces and resist common disinfectant protocols. Some *C. difficile* strains have developed resistance to common antibiotics while also exhibiting more severe pathogenicity (4). Studies estimate that 500,000 CDI cases occur in the U.S. annually (5), resulting in 29,000 deaths and over $4.8 billion in associated costs in acute care facilities alone (6).

Approximately 10% of healthy adults are asymptomatically colonized with *C. difficile* (7-9). Commensal species in the healthy gut can provide resistance against *C. difficile* pathogenic colonization through a variety of metabolic mechanisms, including competition for dietary nutrients such as carbohydrates and amino acids (10) and conversion of host-derived primary bile acids that promote *C. difficile* spore germination to secondary bile acids that inhibit germination and growth (11). Recurrence is a major challenge associated with CDI treatment, as approximately 30% of patients develop a least one occurrence of reinfection (12). The host-microbiota mechanisms underlying recurrence are not well understood, as microbiota composition alone is a poor predictor of patient recovery versus recurrence (13-15). For patients who suffer from repeated episodes of recurrence, fecal microbiota transplantation (FMT) is the last resort treatment. Despite its remarkable success rate approaching 90% (16), FMT remains controversial (17) as the donor microbiota confer poorly understood functions to the endogenous community (18) and may contain pathogenic strains not recognized during screening of donor stool (19).

The advent of metagenomic technologies such as 16S rRNA-encoding gene sequencing has yielded unprecedent insights into the composition of *in vivo* bacterial communities (20-22). Despite numerous metagenomics-based studies that attempt to correlate CDI disease state to bacterial composition (13, 23-26), we still do not understand why some exposed individuals develop CDI while other individuals are asymptomatic (9, 27, 28) and why some infections become recurrent while other infections are effectively treated with antibiotics (29-32). Furthermore, the microbial community being transplanted with FMT is poorly understood both with regard to its composition and the health-promoting metabolic functions being introduced (33-35). Uncertainty at this level can decrease therapeutic efficacy and increase the risk of adverse events (19, 36).

Translating composition data derived from 16S sequencing into an understanding of community function is a challenging problem. Gut bacteria often possess overlapping metabolic functions, such as their ability to synthesize secondary bile acids (37-39) and short-chain fatty acids like butyrate and propionate (40, 41). Furthermore, numerous studies (42-47) have demonstrated that microbiota taxonomic composition is an individual characteristic and usually an inadequate measure for assessing disease states. These critical gaps in knowledge exist because bacterial composition data alone is insufficient to characterize the metabolic state of the diseased gut and nutritional environments that are protective against CDI. The next step in metagenomic applications to microbiome research needs to be the translation of taxa composition data into quantitative information about bacterial community dynamics and function (48-50). In this study, a recently developed *in silico* metagenomics pipeline (mgPipe; 51) was applied to the problem of identifying microbiota-based determinants of recurrent CDI. The pipeline was used to translate 16S-derived taxa abundances from stool samples of CDI and FMT patients into sample-specific models to quantify the metabolic capabilities of the modeled communities, which have been shown to correlate with clinical states in other microbiota-based disease processes (52, 53).

## Materials and Methods

### Patient Data

Gut microbiota composition data were obtained from two published studies (13, 54) in which patient stool samples were subjected to 16S rRNA gene amplicon library sequencing. The first study (13) included 225 longitudinal samples from 93 CDI patients ranging in age from 18 to 89 years. Each patient was characterized as either: *nonrecurrent* if a non-reinfected sample was collected >14 days after a previous *C. difficile* positive sample; *recurrent* if a positive sample was collected 15–56 days after a previous positive sample; and *reinfected* if a positive sample was collected >56 days after a previous positive sample (Table 1). Because patients in both groups were ultimately reinfected, the recurrent and reinfected patients were lumped together in this study and termed recurrent. The sample was defined as an *index* sample if it returned the first *C. difficile* positive for that patient, a *pre-index* sample if it was collected before the index sample, and *post-index* sample if it was collected after the index sample. The second study (54) included 40 samples from 14 FMT patients and 10 of their stool donors (Table 1).

**Table 1.** Summary of CDI patient data (PMC4847246) and FMT patient data (PMC4068257).

| CDI patient data |  |  |  | FMT patient data |  |
| --- | --- | --- | --- | --- | --- |
|  | Nonrecurrent | Recurrent | Total |  | Total |
| Patients | 42 | 51 | 93 | Patients | 14 |
| Pre-index samples | 1 | 5 | 6 | Pre-FMT samples | 14 |
| Index sample | 42 | 51 | 93 | Donor samples | 10 |
| Post-index samples | 37 | 89 | 126 | Post-FMT samples | 26 |
| Total samples | 80 | 145 | 225 | Total samples | 40 |

The 16S rRNA OTU reads available in the two original studies were generally at the genus and family taxonomic levels. These reads were mapped into taxa abundances for development of sample-specific community metabolic models. Using the 100 most abundant OTUs across the samples in each study, taxa abundances were derived as follows: (1) all OTUs belonging to the same taxonomic group were combined; (2) OTUs belonging to higher taxonomic groups (i.e. order and above) were eliminated to maintain modeling at the genus and family levels; and (3) the reduced set of OTUs was normalized such that the abundances of each sample summed to unity. To quantify the effect of eliminating higher-order taxa, the total reads in (3) were divided by the total reads in (2) to generate an unnormalized total abundance for each sample. For the CDI dataset, this procedure resulted in 48 taxa (40 genera, 8 families) that accounted from an average of 97.7% of the top 100 OTU reads across the 225 samples (Table S1). Due to the non-negligible level of the class Gammaproteobacteria in the FMT dataset (average abundance of 3.4%), this class was retained to generate 39 taxa (30 genera, 8 families, 1 class) that accounted for an average of 99.3% of the top 100 OTU reads across the 40 samples (Table S2).

### Community Metabolic Modeling

Taxa represented in the normalized CDI and FMT samples were modeled using genome-scale metabolic reconstructions from the Virtual Metabolic Human (VMH) database (55; www.vmh.life; Figure S1). The function createPanModels within the metagenomics pipeline (mgPipe; 51) of the MATLAB Constraint-Based Reconstruction and Analysis (COBRA) Toolbox (56) was used to create higher taxa models from the 818 strain models available in the VMH database. The sample taxa were mapped to these pan-genome models according to their taxonomy (e.g. Clostridium cluster XI containing *C. difficile* was mapped to the family *Peptostreptococcaceae*). The function initMgPipe was used to construct a community metabolic model for each of the 225 CDI and 40 FMT samples. Model construction required specification of taxa abundances for each sample and maximum uptake rates of dietary nutrients, which was specified according to an average European diet (53; Table S3). The community models contained an average of 33,773 reactions (minimum 8,302; maximum 59,923) for the CDI samples and 36,278 reactions (minimum 26,466; maximum 46,179) for the FMT samples. mgPipe also performed flux variability analysis (FVA) for each model with respect to each of the 411 metabolites assumed to be exchanged between the microbiota and the lumen and fecal compartments. The FVA results were used to compute the net maximal production capability (NMPC; 51) of each metabolite by each model (Table S4 for CDI; Table S5 for FMT) as a measure of community metabolic capability.

### Data analysis

Patient data consisted of normalized taxa abundances and model data consisted of calculated NMPCs, both of which could be connected to associated metadata on a sample-by-sample basis (Tables S1 and S2). Both types of data were subjected to unsupervised learning techniques including kmeans clustering and principal component analysis (PCA) to extract relationships between partitioned samples/patients and clinical parameters such as recurrence. Statistical significance of associations between categorial variables (e.g. recurrent/nonrecurrent) across samples/patient groups were assessed using Fisher’s exact test. Correlations between taxa based on their abundances across samples/patients were calculated using the proportionality coefficient (57), which accounts for the effects of data normalization. Statistically significant differences between metabolite NMPCs across samples/patients were assessed using the Wilcoxon rank-sum test.

## Results

### Clustered index samples were not predictive of recurrence

The 90 index samples remaining after removal of 3 samples containing less than 90% of modeled taxa were clustered using their normalized taxa abundances. The Davies-Bouldin criterion (58) indicated the optimal number of clusters to be 3, and the abundance data were clustered using the kmeans method. The index samples were clustered into 20 samples with elevated *Enterobactericeae* and *Enterococcus*, 33 samples dominated by *Bacteroides*, and 37 samples with elevated *Escherichia* and *Akkermansia* (Figure 1A). While the *Enterobactericeae*/*Enterococcus* cluster exhibited a higher proportion of recurrent samples than the other two clusters and the entire index dataset (Figure 1B), none of these differences were significant (Fisher’s exact test, p > 0.5). When the index samples were analyzed with PCA, the abundance data showed structure with respect to the three clusters but not with respect to recurrent/nonrecurrent samples (Figure 1C).

**Figure 1.**
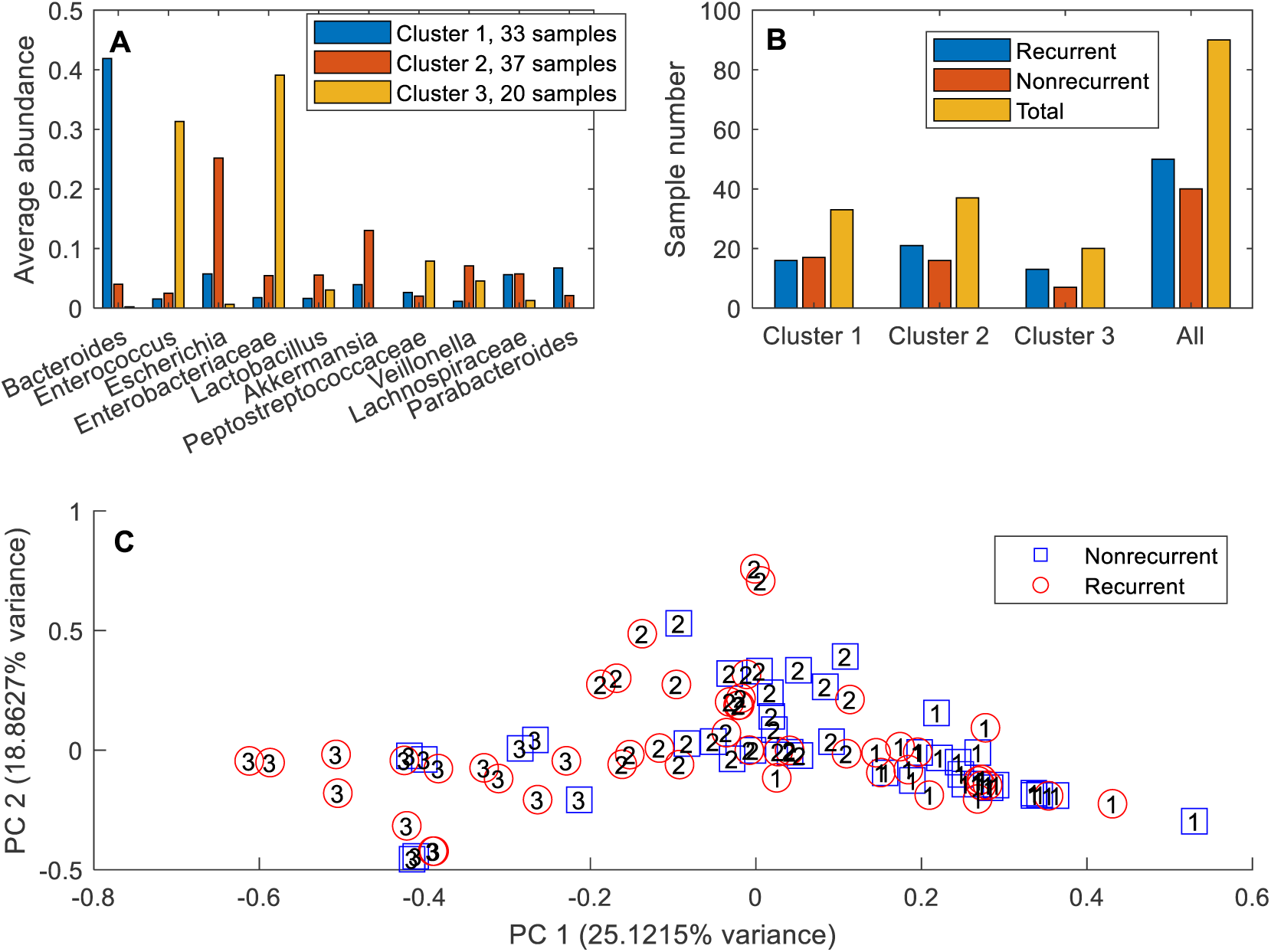
Clustering of 90 index samples using 16S-derived abundance data. (A) Average taxa abundances across the samples in each cluster for taxa which averaged at least 5% of the total abundance. (B) Number of recurrent, nonrecurrent and total samples in each cluster and all 90 index samples. None of the clusters contained a disproportionate number of recurrent samples (Fisher’s exact test, p > 0.5). (C) PCA plot of the abundance data with each recurrent and nonrecurrent sample labeled by its associated cluster number.

Similar analyses were applied to NMPCs calculated for the 90 index samples to determine if the metabolic capability of each community would be more predictive of recurrence than community composition. The index samples were clustered into 25 samples with elevated *Enterobactericeae* and *Escherichia*, 30 samples with elevated *Enterococcus* and *Akkermansia*, and 35 samples dominated by *Bacteroides* (Figure S1A), demonstrating that the abundance data and the model-processed abundance data produced different clustering results (Figure S1B). None of the clusters exhibited a higher proportion of recurrent samples (p > 0.25; Figure S1C), and PCA showed no distinct structure with respect to recurrent/nonrecurrent samples (Figure S1D). Therefore, the index samples, which were collected prior to antibiotic treatment, were deemed to have little predictive value with respect to CDI recurrence.

### Post-index samples clustered by metabolic capability were predictive of recurrence

The 119 post-index samples remaining after removal of samples containing less than 90% of modeled taxa were clustered by their taxa abundances to produce 15 samples dominated by *Enterobactericeae* with low abundances of *Bacteroides, Enterococcus* and *Escherichia*, 18 samples dominated by *Enterococcus* with low abundances of *Bacteroides, Escherichia* and *Enterobactericeae*, and 86 samples more diversely distributed (inverse Simpson index of 12.4 versus 2.2 and 1.9, respectively, for the other two clusters) and not dominated by a single taxa (Figure S2A). The high *Enterobactericeae* cluster contained a disproportionate number of recurrent samples (14/15) compared to the high *Enterococcus* cluster (9/18; Fisher’s exact test, p = 0.009; Figure S2B). The recurrent samples in the high *Enterobactericeae* cluster were clearly distinguishable in a PCA plot of the post-index abundance data (Figure S2C).

NMPCs calculated for the 119 post-index samples were clustered to explore the hypothesis that metabolic outputs of the community models would allow superior recurrence prediction than was possible with community compositions alone. These model-processed sample abundances were clustered into 28 samples with elevated *Enterobactericeae* and *Escherichia*, 28 samples with elevated *Enterococcus* and *Lactobacillus*, and 63 samples with elevated *Bacteroides* and more diversely distributed (inverse Simpson index of 11.6 versus 4.3 and 3.2, respectively, for the other two clusters; Figure 2A). The high *Enterobactericeae* cluster contained a disproportionate number of recurrent samples (25/28) compared to the high *Enterococcus* cluster (14/28; p = 0.003) and the entire post-index dataset (83/119; p = 0.035; Figure 2B). Therefore, the metabolic model generated a larger cluster of high recurrent samples compared to the abundance data (28 samples from 22 patients versus 15 samples from 11 patients) at a higher level of statistical significance. The high recurrence *Enterobactericeae* cluster was distinguishable in the upper left quadrant of a PCA plot of the model-processed abundance data due to the unique metabolic capabilities of these clustered samples (Figure 2C), an issue explored below in detail. Despite having 411 possible PCA components compared to the abundance data with 48 possible components, the model output data was more efficiently compressed with a small number of principal components (e.g. 58.2% versus 48.0% variance captured for 2 components; Figure 2D). Collectively, these results demonstrate the potential benefit of model-based processed abundance data to quantify metabolic functions of sampled communities rather than relying on sample compositions alone.

**Figure 2.**
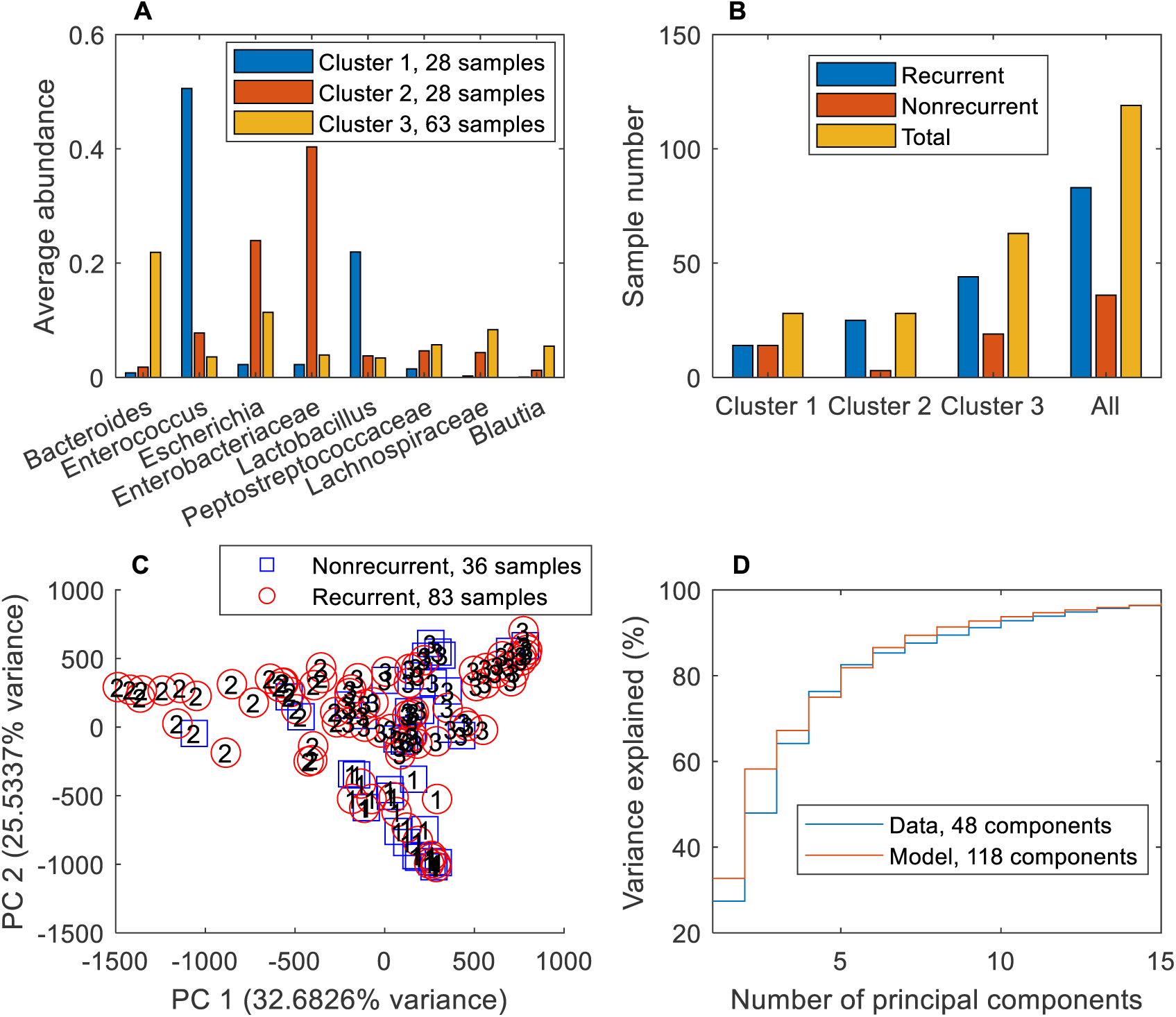
Clustering of 119 post-index samples using model-processed abundance data. (A) Average taxa abundances across the samples in each cluster for taxa which averaged at least 5% of the total abundance. (B) Number of recurrent, nonrecurrent and total samples in each cluster and all 119 post-index samples. Cluster 2 contained a disproportionate number of recurrent samples (25/28) compared to the cluster 1 (14/28; p = 0.003) and the entire post-index dataset (83/119; p = 0.035). (C) PCA plot of model-processed abundance data with each recurrent and nonrecurrent sample labeled by its associated cluster number. (D) Variance explained by PCA of 16S-derived abundance data and model-processed abundance data. The total number of components for each dataset shown in the legend was determined by the MATLAB function pca.

The number of clusters was varied to further explore partitioning of the 119 model-processed post-index samples. Interestingly, 2 clusters also produced a relatively small group with elevated *Enterobactericeae* and *Escherichia* (34 samples) as well as generating a second larger group with elevated *Enterococcus, Bacteroides* and *Lactobacillus* (85 samples; Figure 3A). As the number of clusters was increased, the *Enterobactericeae*/*Escherichia* group split into two separate clusters and the *Enterococcus*/*Bacteroides*/*Lactobacillus* group split into three separate clusters (Figure 3B-E). The high *Enterobactericeae* clusters retained their property of disproportionate recurrence compared to the high *Enterococcus*-elevated clusters for all cases (p < 0.04; Figure 3F), suggesting a possible supportive role for *Enterobactericeae* with respect to CDI recurrence during antibiotic treatment.

**Figure 3.**
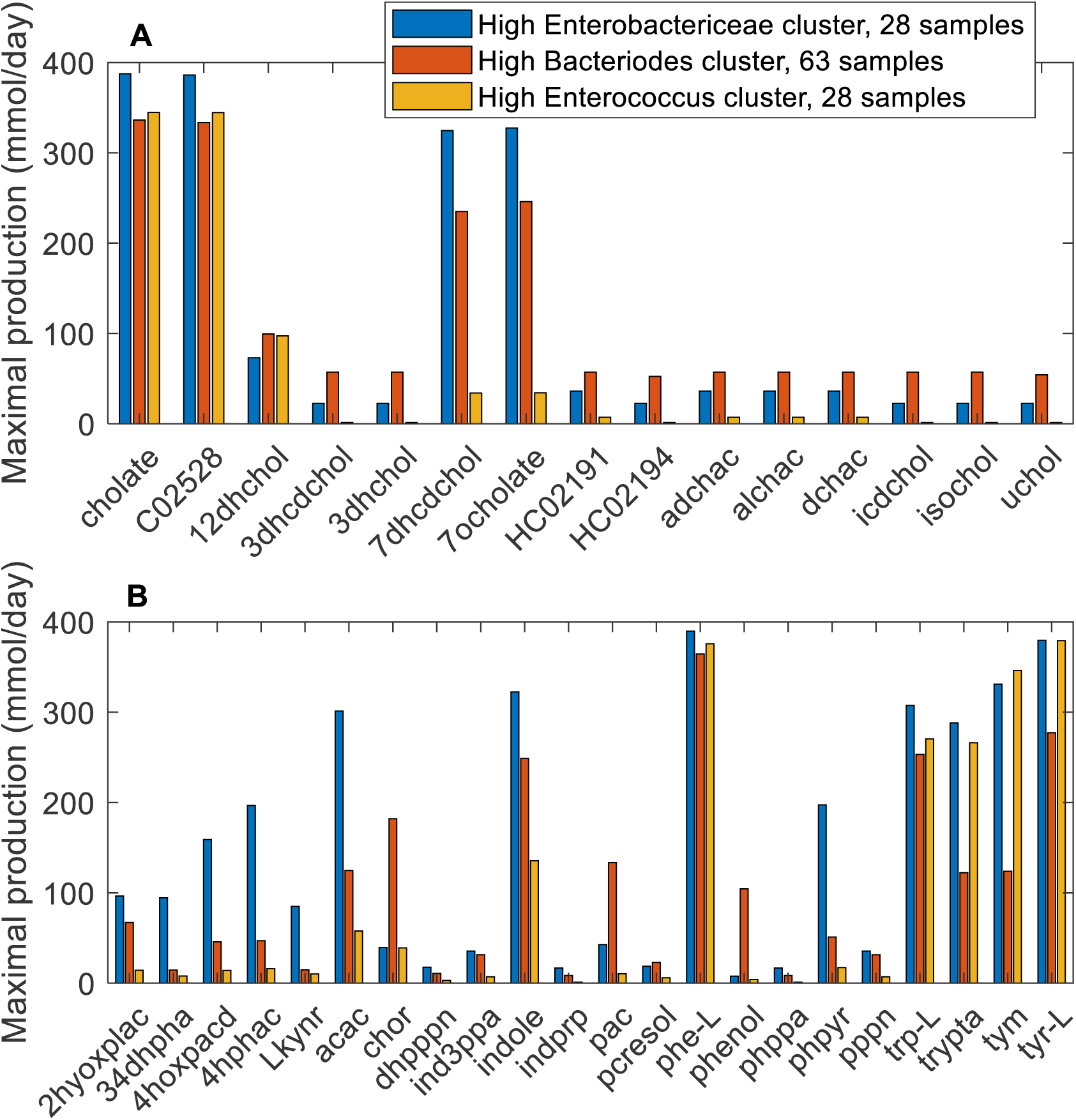
Net maximal production rates of bile acid and aromatic amino acid metabolites in the high *Enterobacteriaceae*, high *Bacteroides* and high *Enterococcus* clusters generated from 119 model-processed post-index samples. (A) Bile acid metabolites in which the average production rate was non-zero in at least one cluster. (B) Aromatic amino acid metabolites in which the average production rate was non-zero in at least one cluster. Metabolites abbreviations are taken from the VMH database (www.vmh.life). Full metabolite names, their associated metabolic pathways and numeric values for their average production rates in each cluster are given in Table S6.

### Clustered post-index samples exhibited distinct bile acid and aromatic amino acid metabolism

NMPCs of the 119 post-index samples with respect to each of the 411 exchanged metabolites were statistically analyzed to assess metabolic differences between the 3 clusters. For each pair of clustered samples, the Wilcoxon rank sum test was applied to the NMPCs on a metabolite-by-metabolite basis. To reduce the number of reported metabolites, statistically different metabolite NMPCs (p < 0.05) also were required to have an average NMPC > 50 mmol/day in at least one cluster and average NMPC that differed between the clusters by at least 100%. A comparison of the high *Enterobactericeae* cluster (HEb, 28 samples) and the high *Enterococcus* cluster (HEc, 28 samples) generated 44 differentially produced metabolites (Figure S5, Table S6), with 19 metabolites associated with aromatic amino acid (AAA), bile acid (BA) and butanoate metabolism. The HEb cluster and the high *Bacteroides* cluster (HBo, 63 samples) had 47 differentially produced metabolites (Figure S6, Table S6), including 7 metabolites associated with AAA degradation elevated in the HEb cluster. Interestingly, 11 secondary BA metabolites were elevated in the HEc cluster compared to the HBo cluster, accounting for 25% of the differentially produced metabolites (Figure S7, Table S6).

Due to their differential utilization across the 3 clusters, the BA and AAA pathways were examined more carefully by collecting all metabolites belonging to these pathways that were allowed to be exchanged according to the metabolic models. The HEb cluster had the highest production capabilities for the two unconjugated primary BAs (Figure 3A), which have been reported to either promote (cholate) or inhibit (chenodeoxyholate, C02528) *C. difficile* germination (59, 60). By contrast, the HBo cluster generated the highest production of most secondary BAs, which are known to be generally protective against CDI (2, 61, 62). Interestingly, the HEc cluster had much lower production capabilities for secondary BAs. The HEb cluster consistently generated higher production of metabolites involved in AAA catabolism but not significantly higher production of the AAAs themselves (Figure 3B, Table S6). This predicted AAA degradation ability was decreased in the HBo cluster and substantially lower in the HEc cluster, with the notable exceptions of the tyrosine degradation product tyramine (tyr) and the tryptophan-derived metabolite tryptamine (trypta). Interestingly, the key AAA precursor chorismite (chor) was significantly elevated in the HEc cluster, yet the production capabilities of the AAA themselves were reduced in this cluster. Since the HEb cluster contained a disproportionate number of recurrent samples compared to the other 2 clusters, these predictions suggest a possible role for AAA metabolism in CDI recurrence.

### Transient presence in the high Enterobactericeae cluster was sufficient for elevated patient recurrence

The HEb cluster contained a disproportionate number of recurrent samples (25/28). To investigate the transient bacterial communities of the 22 patients from which these samples were collected, all post-index samples from these patients were grouped to generate an enlarged dataset of 55 samples. Similarly, all post-index samples from the 46 patients in the HBo cluster and the 21 patients in the HEc cluster were grouped to generates datasets containing 87 and 47 samples, respectively. The 66 total patients represented by these samples were allowed to reside in multiple datasets referred to as the HEb, HBo and HEc groups. The HEb group contained a disproportionate number of recurrent patients (19/22) compared to the HEc group (12/21; p = 0.045) and all grouped patients (41/66; p = 0.038; Figure 4A). Within the HEb group, *Enterobacteriaceae* was most negatively correlated with *Escherichia* and *Bacteroides* (proportionality coefficient ρ = −0.18 for both pairs; Figure 4B). In addition to health-promoting *Bacteroides* (63), *Enterobacteriaceae* was negatively correlated with other taxa including *Lachnospiraceae* (64), *Lactobacillus* (65) *Akkermansia* (66) and *Alistipes* (65) reported to be protective against CDI.

**Figure 4.**
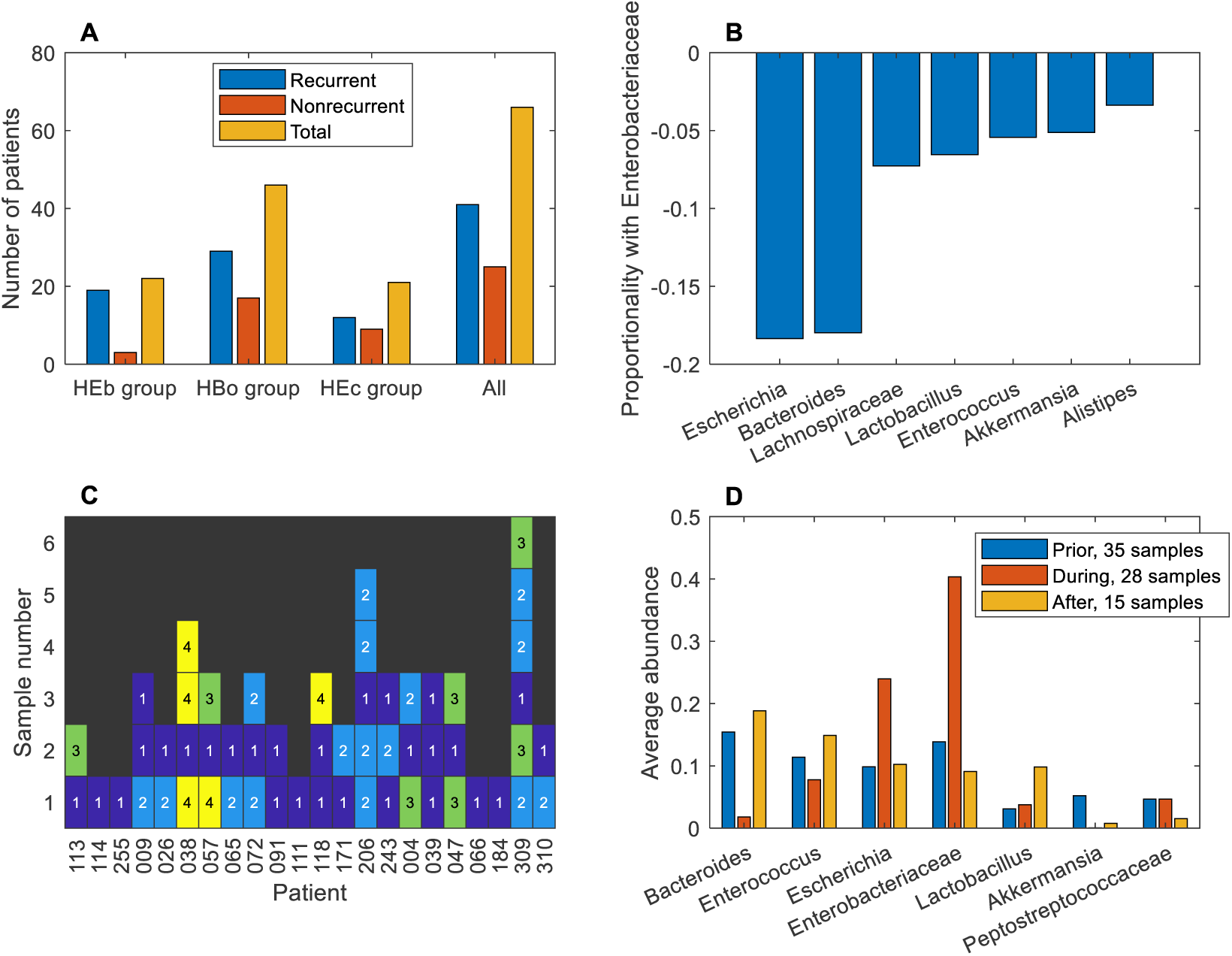
Analysis of patient samples in the high *Enterobacteriaceae* (HEb) group. All post-index samples from the 22 patients with at least one sample in the high *Enterobacteriaceae* (HEb) cluster were grouped to generate the enlarged HEb group of 55 samples. (A) The number of recurrent, nonrecurrent and total patients in the HEb group compared to those in the HBo and HEc groups. All post-index samples of the 46 patients represented in the HBo cluster and the 21 patients represented in the HEc cluster were grouped to produce the 87 and 47 samples, respectively, in the HBo and HEc groups. The 66 total patients represented by these samples were allowed to reside in multiple groups. (B) Correlation between *Enterobacteriaceae* and other taxa in the HEb group calculated from the 55 post-index samples as measured by the proportionality coefficient ρ (PMC4361748). The 7 taxa with the largest |ρ| values are shown. (C) Transient progression of samples from the 22 patients in the HEB group with samples denoted as 1 if contained in the HEb cluster, 2 if contained in the HBo cluster, 3 if contained in the HEc cluster and 4 if not clustered due to low abundance of modeled taxa (see Methods). (D) Average taxa abundances for an expanded HEb group that also contained pre-index and index samples to generate a dataset of 78 samples. These samples were partitioned into 35 samples prior to patients entering the HEb cluster, 28 samples during patient presence in the HEb cluster and 15 samples after patients left the HEb cluster.

Taxa abundances from the HEb group showed considerable dissimilarity between samples from an individual patient (Yue and Clayton dissimilarity index θ = 0.35 across the 22 patients; Figure S8), with the only consistent feature among the 22 patients being at least one sample contained in the HEb cluster (Figure 4C). The first 3 patients (113, 114, 255) were nonrecurrent despite having a sample in the HEb cluster. Of the remaining 19 recurrent patients, 10 patients also had a sample in the HBo cluster and 4 patients also had a sample in the HEc cluster. Moreover, 7 of the 19 recurrent patients had final samples contained in either the HBo or HEc cluster. Given the irregular sampling frequency reported in the original clinical study (13), this analysis suggests that transient presence in the HEb cluster was sufficient to have an elevated risk of recurrence.

To investigate if the bacterial community within an individual patient showed distinct trends once the HEb cluster was entered, the HEb group was expanded to contain index samples and partitioned into 35 samples prior to patients entering the HEb cluster, 28 samples during patient presence in the HEb cluster and 15 samples after patients left the HEb cluster. For each patient, the Yue and Clayton dissimilarity index θ was calculated using taxa abundances in the last sample before entering the HEb cluster, the first sample in the cluster and the first sample after leaving the cluster (if such a sample existed). Samples showed more dissimilarity when leaving the cluster (θ = 0.11; Figure 4C) than when entering the cluster (θ = 0.27). Interestingly, samples entering and leaving the clusters were the most similar (θ = 0.35), suggesting partial community restoration following transient presence in the cluster. Within the HEb group, the only significant taxa abundance changes upon entering the HEb cluster were a large drop in *Bacteroides* (Wilcoxon rank sum test, p < 0.006) and the expected large increase in *Enterobacteriaceae* (p < 0.004). The abundances of these taxa subsequently returned to near pre-HEb values upon leaving the cluster. Collectively, these analyses suggest a possible role for *Bacteroides* in opposing recurrent infection.

Transient presence in the HEb cluster was not associated with a concurrent or future increase in the abundance of *Peptostreptococcaceae* (Figure 4D), the family containing *C. difficile*. In fact, *Enterobacteriaceae* and *Peptostreptococcaceae* abundances were only weakly correlated within the entire HEb group (ρ = −0.01). Therefore, transient presence in the HEb cluster was hypothesized to temporarily create a metabolic environment that promoted CDI recurrence through an increase in *C. difficile* toxicity rather than *C. difficile* expansion. To explore this hypothesis, the metabolite production capabilities of the HEb, HBo and HEc groups were compared. The metabolic signature of the HEb group (Figure S9) was similar to that predicted when the HEb and HEc clusters were compared (Figure S6, Table S6) and included elevated synthesis of metabolites known to induce (e.g. butyrate) and suppress (e.g. cysteine) the toxicity of *C. difficile* (67, 68).

### FMT patient samples clustered by metabolic capability were predictive of sample type

Taxa abundance data derived from 40 stool samples representing 14 recurrent CDI patients undergoing FMT and from their donors (54) were modeled and to investigate the community metabolic changes taking place upon FMT treatment. Model-processed abundance data were clustered to generate one small cluster with elevated *Cronobacter, Enterobacteriaceae* and *Gammaproteobacteria* (averaged 67.9% across the 11 samples) and a second larger cluster with elevated *Bacteroides* and *Lachnospiraceae* (averaged 43.6% across the 29 samples; Figure 5A). Since *Cronobacter* belongs to the family *Enterobacteriaceae* and these two taxa averaged 56.4% across the 11 samples, the small cluster was considered to be dominated by *Enterobacteriaceae*. Consistent with these results, *Cronobacter* was most positively correlated with *Enterobacteriaceae* (proportionality coefficient ρ = +0.12) but negatively correlated with *Bacteroides* (ρ = −0.29) and several other taxa (*e*.*g. Lachnospiraceae*; Figure 5B) often reported to be CDI protective (64, 65). Similarly, *Bacteroides* was negatively correlated with *Enterobacteriaceae* (ρ = −0.21), *Gammaproteobacteria* and *Clostridiaceae* (Figure S10B), taxa which have been reported to be elevated in CDI (64, 65, 69).

**Figure 5.**
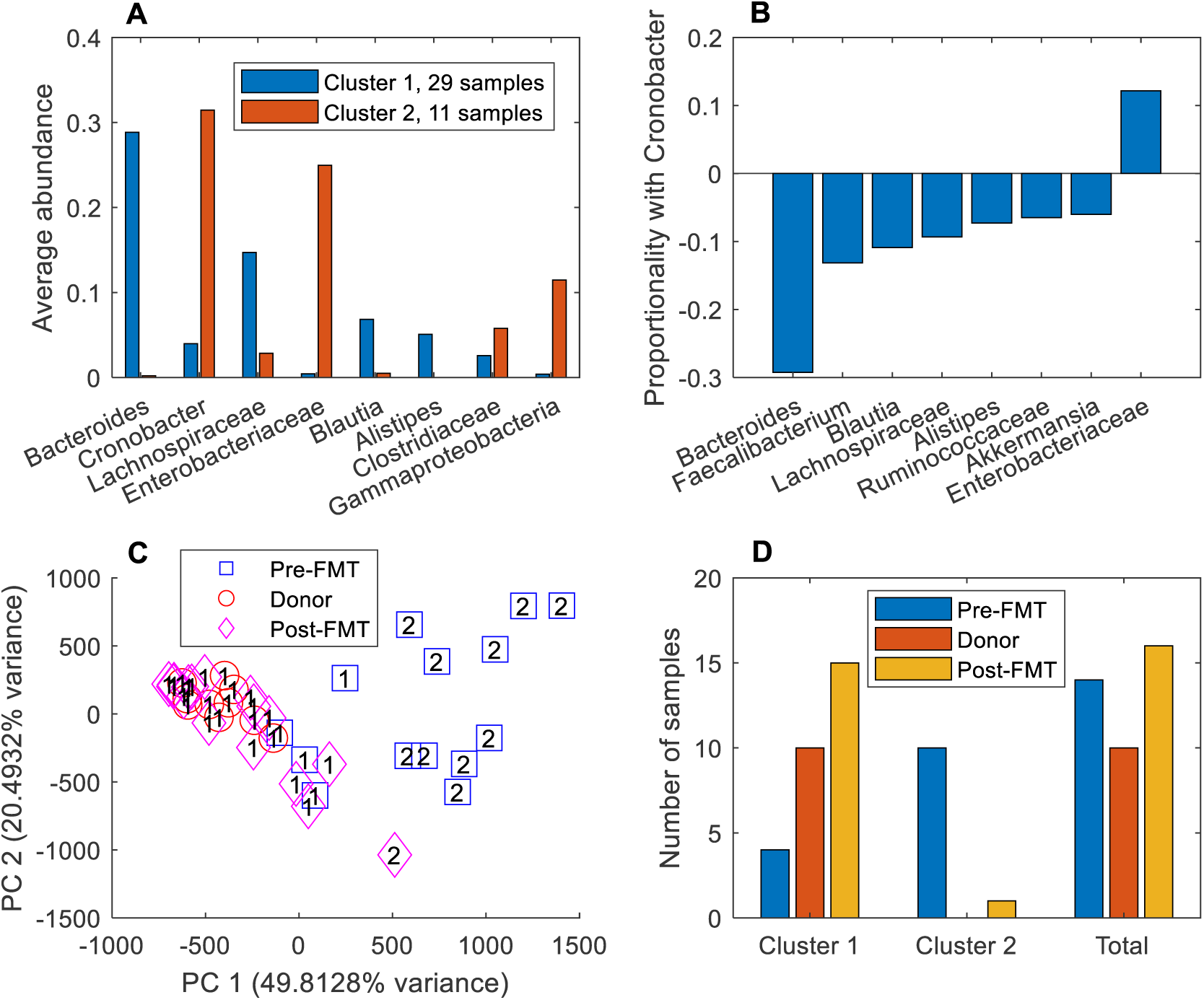
Clustering of 40 FMT samples using model-processed abundance data. (A) Average taxa abundances across the samples in each cluster for taxa which averaged at least 5% in at least one cluster. (B) Correlations between *Cronobacter* and other taxa calculated from all 40 samples as measured by the proportionality coefficient p. The 8 taxa with the largest |ρ| values are shown. (C) PCA plot of the model-processed abundance data with each pre-FMT, donor and post-FMT sample labeled by its associated cluster number. (D) Number of pre-FMT, donor and post-FMT samples in each cluster and all 40 samples. Cluster 2 contained a disproportionately large number of pre-FMT samples (10/11) compared to the cluster 1 (4/29; p < 0.0001) and the entire FMT dataset (14/40; p = 0.0014).

When PCA was performed on the model-processed abundance data, the *Enterobacteriaceae*-dominated cluster was clearly distinguishable and appeared to have an overrepresentation of pre-FMT patient samples (Figure 5C). In fact, this cluster contained a disproportionately large number of pre-FMT samples (10/11) compared to both the *Bacteroides*-elevated cluster (4/29; p < 0.0001) and the entire sample set (14/40; p = 0.0014; Figure 5D). Additionally, the *Enterobacteriaceae*-dominated cluster had a disproportionately small number of donor samples (0/11) and post-FMT patient samples (1/11) compared to the *Bacteroides*-elevated cluster (p = 0.038 and 0.027, respectively). The findings that the high *Enterobacteriaceae* (HEb) cluster studied earlier contained a disproportionately large number of recurrent CDI samples (see Figure 2) and the high *Enterobacteriaceae* cluster found here contained a disproportionately large number of pre-FMT samples provide additional support for the hypothesis that elevated *Enterobacteriaceae* is associated with recurrent CDI.

When similar analyses were applied directly to the abundance data, the dataset was split equally into one cluster with elevated *Cronobacter* and *Enterobacteriaceae* (averaged 35.9% across the 20 samples) and a second cluster with elevated *Bacteroides* and *Lachnospiraceae* (averaged 55.5% across the 20 samples; Figure S10A,C). The *Enterobacteriaceae*-elevated cluster contained all the pre-FMT samples (14/20), representing large statistical differences with the *Bacteroides*-elevated cluster (0/20; p < 0.00001) and the entire dataset (14/40; p = 0.0014; Figure S10D). By contrast, the *Bacteroides*-elevated cluster contained a disproportionally large number of donor samples (9/20) compared to the *Enterobacteriaceae*-elevated cluster (1/20). These results are consistent with those obtained from the model-processed abundance data and collectively identified the pre-FMT samples as compositionally and functionally distinct from the donor and post-FMT samples.

### High Enterobactericeae FMT cluster exhibited distinct bile acid and aromatic amino acid metabolism

Predicted NMPCs of 411 exchanged metabolites were statistically analyzed to assess metabolic differences between the 40 samples clustered based on model-processed abundance data. A comparison of the high *Enterobactericeae* cluster (HEb-FMT, 11 samples) and the high *Bacteroides* cluster (HBo-FMT, 29 samples) generated 60 differentially produced metabolites (Figure S11). Only 23 of these 60 metabolites were identified as being differentially produced between the HEb and HBo clusters defined from model processing of CDI post-index samples (Figure S5; Table S6). Interestingly, 10 secondary BA metabolites and 4 AAA catabolic products were among the 37 newly identified metabolites. Therefore, BA and AAA metabolism in the HEb-FMT and HBo-FMT clusters were examined more carefully by comparing all secreted metabolites belonging to these pathways. The HEb-FMT cluster had decreased production of all 17 BA metabolites (Figure 6A), including significantly reduced synthesis of 10 secondary BAs generally correlated with recurrent CDI (59, 70, 71). By contrast, the HEb-FMT cluster had enhanced AAA metabolism as evidenced by elevated production of all 3 AAAs and 15 AAA degradation products, including significantly increased synthesis of 8 degradation products (Figure 6B). Given that the HEb-FMT cluster was overrepresented in pre-FMT samples and underrepresented in donor and post-FMT samples, these predictions provide additional support for the hypothesis that BA and AAA community metabolism may play key roles in CDI recurrence and treatment.

**Figure 6.**
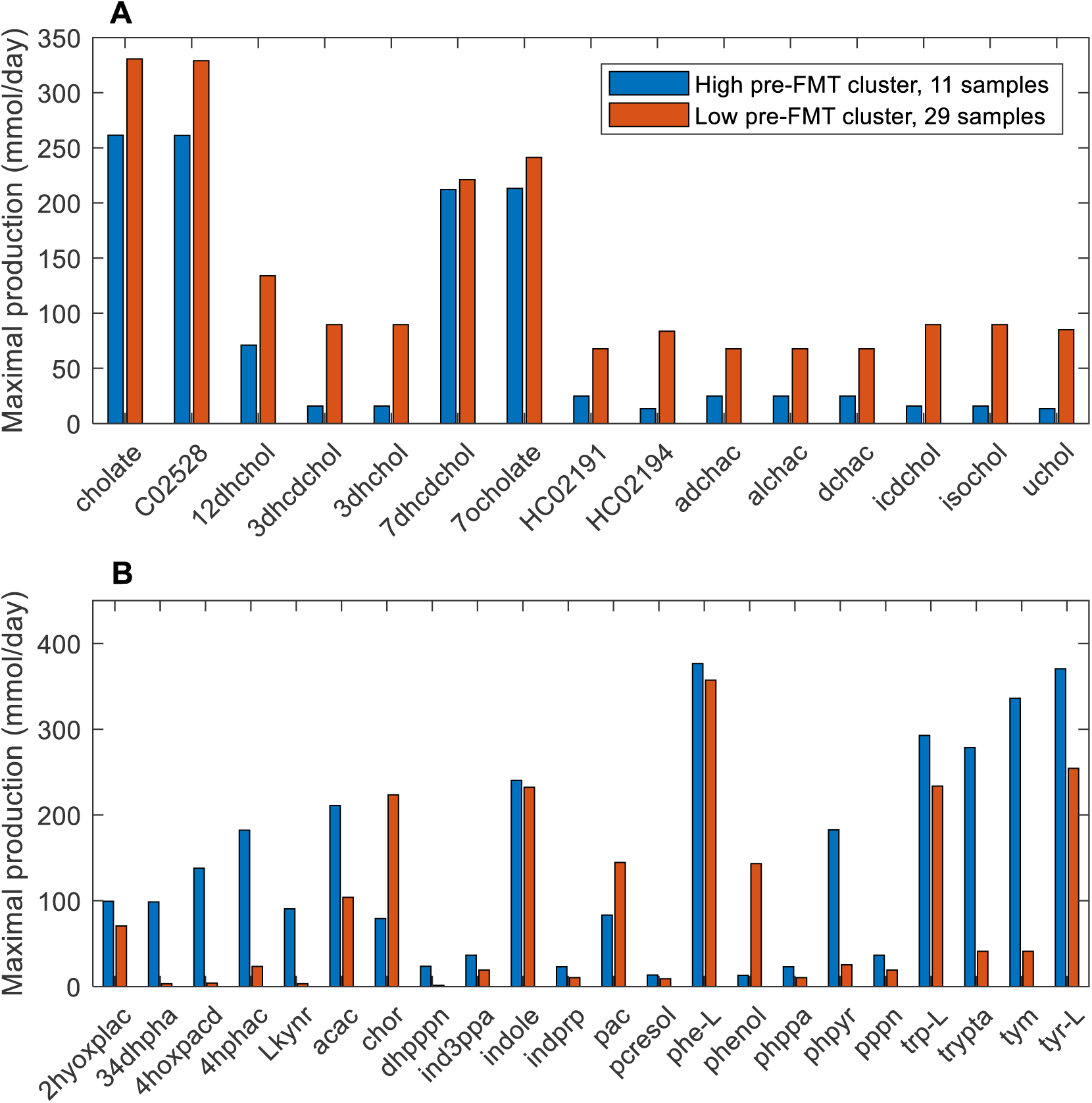
Net maximal production rates of bile acid and aromatic amino acid metabolites in the high *Enterobacteriaceae* and high *Bacteroides* clusters generated from 40 model-processed FMT samples. (A) Bile acid metabolites in which the average production rate was non-zero in at least one cluster. (B) Aromatic amino acid metabolites in which the average production rate was non-zero in at least one cluster. Metabolite abbreviations are taken from the VMH database (www.vmh.life). Full metabolite names, their associated metabolic pathways and numeric values for their average production rates in each cluster are given in Table S7.

When the same analysis procedure was applied to NMPCs clustered according to 16S-derived abundance data, 46 metabolites differentially produced between the *Enterobactericeae*-elevated and *Bacteroides*-elevated clusters were identified (Figure S12). Overproduction of AAA catabolic products in the *Enterobactericeae*-elevated cluster continued to be pronounced, but differences in secondary BAs between the two clusters were no longer evident. The inability of the clustered abundance data to generate differential predictions of BA metabolism was attributed to the *Enterobactericeae*-elevated cluster containing 1 donor and 5 post-FMT samples in addition to all 14 pre-FMT samples. Therefore, clustering the samples according to model-processed abundance data appeared to offer advantages for understanding community metabolic changes resulting from FMT.

## Discussion

An *in silico* metagenomics pipeline was used to translate 16S-derived abundance data into sample-specific community models for investigating the metabolic determinants of recurrent CDI. The models allowed sample-by-sample predictions of metabolite production rates that were used both to cluster samples according to their functional metabolic capabilities and to provide mechanistic insights into clusters exhibiting high recurrence. Community model predictions were dependent both on the taxonomic groups represented in the 16S data and the fidelity of individual taxa metabolic models. The CDI (PMC4847246) and FMT (54) datasets used in this study captured taxonomic differences primary at the genus and family levels and therefore precluded modeling metabolism at the strain and species levels (53). Despite this limitation, the pan-genome metabolic models used for community modeling allowed substantial differentiation of samples according to their functional capabilities.

Taxa abundance data and model-processed abundance data were clustered to determine if the resulting clusters exhibited statistically significant differences between the number of recurrent CDI samples. No significant differences were observed when only index samples were tested, suggesting that community composition prior to CDI treatment may provide limited information about recurrence. By contrast, both abundance data and model-processed abundance data derived from post-index samples identified high *Enterobacteriaceae*, low *Bacteroides* clusters as having disproportionate numbers of recurrent samples. Numerous studies have identified *Enterobacteriaceae* as positively associated and *Bacteroides* as negatively associated with primary CDI (63-65) and to a lesser extent with subsequent reinfection (72, 73). The analyses presented here suggest CDI recurrence is more dependent on community response to antibiotic therapy than on the community composition entering therapy. Indeed, first-line antibiotics for CDI treatment including metronidazole and vancomycin are known to collaterally target *Bacteroides* (74, 75) while having little efficacy against *Enterobacteriaceae* (76-78). Unfortunately, the metadata available for these samples only reported if the patient received antibiotic therapy prior to CDI diagnosis. Since next generation antibiotics such as fidaxomicin used for recurrent CDI are more specific for *C. difficile* and are known to spare *Bacteroides* (79, 80), knowledge of which antibiotics were used to treat the patients represented in the high *Enterobacteriaceae* clusters would enable additional analysis.

As compared to direct use of abundance data, an advantage of utilizing predicted metabolite production rates for sample clustering was that the high *Enterobacteriaceae* (HEb) cluster contained more samples (28 vs. 15) representing more patients (22 vs. 11). The model-based cluster included samples with a high combination of *Enterobacteriaceae* and *Escherichia*, which have similar metabolic capabilities since *Escherichia* is a genus within *Enterobacteriaceae*. The capability to collapse samples with different compositions but similar metabolic features is useful when dealing with 16S-derived abundance data at several taxonomic levels, a common situation in human microbiome research.

Another benefit of quantifying metabolic capabilities through modeling was the ability to predict differentially synthesized metabolites across sample groups. When compared to a more taxonomically diverse cluster with elevated *Bacteroides* (HBo cluster) and no statistical difference in recurrence, the HEb cluster was predicted to have significantly reduced capabilities for secondary bile acid (BA) synthesis. These predictions were generally consistent with the established role of BA metabolism in recurrent CDI, as elevated primary BA and reduced secondary BA levels are known to be a disease signature (59, 70, 71). The specific effects of individual BA metabolites are more nuanced, as the primary BA cholate is known to induce germination of *C. difficile* spores, the primary BA chenodeoxycholate suppresses both germination and vegetative growth and the secondary BA deoxycholate induces germination but suppresses growth (64, 81). To achieve prediction at this level of granularity, the metabolic models would need to be constructed with 16S-derived abundance data at lower taxonomic levels since individual species and strains are known to have distinct BA metabolism (53).

Despite having no statistical difference in recurrence, a third cluster elevated in *Enterococcus* and to a lesser extent *Lactobacillus* (HEc cluster) had significantly reduced capabilities for secondary BA synthesis compared to both the HEb and HBo clusters. These predictions underscore the fact that recurrent CDI is a complex disease and not likely to be completely explained by a single factor such as community BA metabolism (54, 65). Interestingly, model-based analysis revealed aromatic amino acid (AAA) metabolism as a second putative mechanism underlying increased recurrence in the HEb cluster. More specifically, this cluster was predicted to have significantly increased synthesis of numerous AAA degradation products compared to the two lower recurrence clusters. *Enterobacteriaceae* is thought to be largely responsible for AAA catabolism in the gut (41, 82), and AAA synthesis has been implicated as a metabolic function protective against CDI (83). *C. difficile* isolates have been shown to have highly variable AAA metabolisms (84), opening the possibility that *Enterobacteriaceae* interactions with *C. difficile* are isolate dependent. However, the 22 patients represented in the HEb cluster were reported to have been infected with at least 9 different *C. difficile* ribotypes. While evidence directly linking AAA metabolism and CDI is currently lacking, the modeling work presented here suggests that this putative connection could be a fruitful area for future experimental studies.

All samples from each patient with at least one sample in the HEb cluster were collected to allow longitudinal analysis of individual patients. This HEb group had a disproportionate number of recurrent patients (19/22) compared to the larger patient population. HEb group patients exhibited compositionally variable communities that routinely switched between clusters, suggesting that transient presence in the HEb cluster could be sufficient for CDI recurrence. Since *Enterobacteriaceae* and *C. difficile* abundances had a very weak negative correlation within the HEb group, *Enterobacteriaceae* did not seem to support *C. difficile* vegetative growth but may have induced spore germination and/or enhanced toxicity of vegetative cells. As discussed above, the BA metabolite profile predicted for the HEb cluster was consistent with enhanced germination. *C. difficile* toxicity is thought to be regulated by a number of metabolites (67, 68, 85). Two of the most potent regulators are toxicity-inducing butyrate and toxicity-suppressing cysteine, both of which were predicted to be elevated in the HEb cluster so as to have opposing effects. An intriguing but entirely speculative possibility is that AAA degradation products from *Enterobacteriaceae* induced *C. difficile* toxicity.

To test consistency of model predictions derived from the CDI dataset, the *in silico* metagenomics pipeline was applied to 40 samples obtained from FMT patients and their stool donors (54). Clustering of model-processed abundance data generated a cluster with a disproportionate number of pre-FMT samples, suggesting distinct metabolic function compared to donor and post-FMT communities as has been reported (70, 86, 87). This cluster had elevated *Cronobacter* and *Enterobacteriaceae* with very low *Bacteroides*. Because *Cronobacter* is a member of *Enterobacteriaceae*, this cluster was identified as high *Enterobacteriaceae* and was compositionally similar with the high recurrence HEb cluster found in the CDI dataset. A second cluster comprised mainly of donor and post-FMT samples was elevated in *Bacteroides* and *Lachnospiraceae* and compositionally similar to the HBo cluster identified from CDI samples. Consistent with these results, *Cronobacter* was found to be strongly positively correlated with *Enterobacteriaceae* and strongly negatively correlated with *Bacteroides* across the FMT dataset. These predictions agreed with observations that FMT tends to decrease the abundances of *Enterobacteriaceae* and other Proteobacteria (54, 73, 88) while increasing the abundances of *Bacteroides* and other health-promoting taxa such as *Lachnospiraceae, Blautia* and *Alistipes* (35, 89, 90).

The HEb cluster identified from FMT samples (HEb-FMT) was predicted to have reduced capabilities for synthesis of both primary and secondary BAs, while the HEb cluster derived from CDI samples (HEb-CDI) exhibited only reduced secondary BA synthesis. Unlike the pan-genome model of the family *Enterobacteriaceae*, the *Cronobacter* genus model lacked BA metabolic pathways because the necessary deconjugation and transformation genes have not been identified in *Cronobacter sakazakii*, the only member of this genus contained in the VMH database. The predicted difference in primary BA synthesis capabilities between *Enterobacteriaceae* in the CDI samples and *Cronobacter*/*Enterobacteriaceae* in the FMT samples demonstrate possible limitations of metabolic modeling at higher taxonomic levels and the potential value of more resolved 16S rRNA sequence data. Despite these differences, the HEb-FMT cluster still exhibited reduced secondary BA levels observed in recurrent CDI (59, 70, 71) and resolved through FMT (70, 71, 91). The HEb-FMT cluster also was predicted to have the capability for elevated AAA degradation including increased synthesis of the catabolic products phenylpyruvic acid, tyramine and tryptamine derived from phenylalanine, tryrosine and tryptophan, respectively. Because they also were elevated in the HEb-CDI cluster compared to high Bacteroides (HBo-CDI) cluster, these 3 metabolites might make interesting experimental targets for their ability to induce germination and/or enhance toxicity of *C. difficile* clinical isolates.

Despite the ability of the proposed *in silico* workflow to identify high *Enterobacteriaceae*-containing communities as disproportionally recurrent and pre-FMT, model-based clustering did not result in idealized partitioning of patient samples. For example, the HEb-CDI cluster contained 3 nonrecurrent patients along with 19 recurrent patients, and the HEb-FMT cluster contained 1 post-FMT sample along with 10 pre-FMT samples. Similarly, the HEb clusters contained only subsets of all recurrent patients (22/66) and all pre-FMT samples (10/14). One possible explanation was that the likelihood of recurrence was dependent on the duration the transient community had an HEb cluster-like composition, as *Enterobacteriaceae* would require sufficient time to establish favorable metabolic conditions for *C. difficile* pathogenicity. While intriguing, such speculation was impossible to test with the available dataset due to infrequent and irregular sampling. The most obvious explanation for the observed discrepancies is that recurrent CDI has a very complex disease etiology that depends on host-microbiota-environment interactions, both metabolic and non-metabolic. Therefore, the inability to fully predict patient recurrence based only on model-processed 16S-derived abundance was hardly surprising. However, the hypotheses that high *Enterobacteriaceae*-containing communities are more prone to recurrence and that recurrence may be partially attributable to the combination of disrupted BA and AAA metabolism seems worthy of further investigation through the type of integrated metagenomics-modeling framework utilized in this study.

## Acknowledgements

The author wishes to acknowledge his Ph.D. student Ayushi Patel for her assistance with generating the references.

## References

1. Surawicz CM, Brandt LJ, Binion DG, Ananthakrishnan AN, Curry SR, Gilligan PH, McFarland LV, Mellow M, Zuckerbraun BS. 2013. Guidelines for diagnosis, treatment, and prevention of Clostridium difficile infections. The American journal of gastroenterology 108:478.

2. Pérez-Cobas AE, Moya A, Gosalbes MJ, Latorre A. 2015. Colonization resistance of the gut microbiota against Clostridium difficile. Antibiotics 4:337–357.

3. Theriot CM, Young VB. 2015. Interactions between the gastrointestinal microbiome and Clostridium difficile. Annu Rev Microbiol 69:445–461.

4. Jarrad AM, Karoli T, Blaskovich MA, Lyras D, Cooper MA. 2015. Clostridium difficile drug pipeline: challenges in discovery and development of new agents. J Med Chem 58:5164–5185.

5. Lessa FC, Mu Y, Bamberg WM, Beldavs ZG, Dumyati GK, Dunn JR, Farley MM, Holzbauer SM, Meek JI, Phipps EC. 2015. Burden of Clostridium difficile infection in the United States. New Engl J Med 372:825–834.

6. Dubberke ER, Olsen MA. 2012. Burden of Clostridium difficile on the healthcare system. Clin Infect Dis 55:S88–S92.

7. Ozaki E, Kato H, Kita H, Karasawa T, Maegawa T, Koino Y, Matsumoto K, Takada T, Nomoto K, Tanaka R. 2004. Clostridium difficile colonization in healthy adults: transient colonization and correlation with enterococcal colonization. J Med Microbiol 53:167–172.

8. Poutanen SM, Simor AE. 2004. Clostridium difficile-associated diarrhea in adults. Cmaj 171:51–58.

9. Furuya-Kanamori L, Marquess J, Yakob L, Riley TV, Paterson DL, Foster NF, Huber CA, Clements AC. 2015. Asymptomatic Clostridium difficile colonization: epidemiology and clinical implications. BMC Infect Dis 15:516.

10. Chang JY, Antonopoulos DA, Kalra A, Tonelli A, Khalife WT, Schmidt TM, Young VB. 2008. Decreased diversity of the fecal microbiome in recurrent Clostridium difficile—associated diarrhea. The Journal of infectious diseases 197:435–438.

11. Sorg JA, Sonenshein AL. 2010. Inhibiting the initiation of Clostridium difficile spore germination using analogs of chenodeoxycholic acid, a bile acid. J Bacteriol 192:4983–4990.

12. Dubberke E. 2012. Clostridium difficile infection: the scope of the problem. Journal of hospital medicine 7:S1–S4.

13. Seekatz AM, Rao K, Santhosh K, Young VB. 2016. Dynamics of the fecal microbiome in patients with recurrent and nonrecurrent Clostridium difficile infection. Genome medicine 8:47.

14. Schubert AM, Rogers MA, Ring C, Mogle J, Petrosino JP, Young VB, Aronoff DM, Schloss PD. 2014. Microbiome data distinguish patients with Clostridium difficile infection and non-C. difficile-associated diarrhea from healthy controls. MBio 5:e01021–14.

15. Antharam VC, Li EC, Ishmael A, Sharma A, Mai V, Rand KH, Wang GP. 2013. Intestinal dysbiosis and depletion of butyrogenic bacteria in Clostridium difficile infection and nosocomial diarrhea. J Clin Microbiol 51:2884–2892.

16. Austin M, Mellow M, Tierney WM. 2014. Fecal microbiota transplantation in the treatment of Clostridium difficile infections. The American journal of medicine 127:479–483.

17. Tan X, Johnson S. 2019. Fecal microbiota transplantation (FMT) for C. difficile infection, just say ‘No’. Anaerobe:102092.

18. Chang C-S, Kao C-Y. 2019. Current understanding of the gut microbiota shaping mechanisms. J Biomed Sci 26:1–11.

19. Wang S, Xu M, Wang W, Cao X, Piao M, Khan S, Yan F, Cao H, Wang B. 2016. Systematic review: adverse events of fecal microbiota transplantation. PloS one 11:e0161174.

20. Costea PI, Hildebrand F, Arumugam M, Bäckhed F, Blaser MJ, Bushman FD, De Vos WM, Ehrlich SD, Fraser CM, Hattori M. 2018. Enterotypes in the landscape of gut microbial community composition. Nature microbiology 3:8–16.

21. Hugerth LW, Andersson AF. 2017. Analysing microbial community composition through amplicon sequencing: from sampling to hypothesis testing. Frontiers in Microbiology 8:1561.

22. Ashton JJ, Beattie RM, Ennis S, Cleary DW. 2016. Analysis and interpretation of the human microbiome. Inflammatory bowel diseases 22:1713–1722.

23. Zhang L, Dong D, Jiang C, Li Z, Wang X, Peng Y. 2015. Insight into alteration of gut microbiota in Clostridium difficile infection and asymptomatic C. difficile colonization. Anaerobe 34:1–7.

24. Leslie JL, Vendrov KC, Jenior ML, Young VB. 2019. The gut microbiota is associated with clearance of Clostridium difficile infection independent of adaptive immunity. mSphere 4:e00698–18.

25. Amrane S, Hocquart M, Afouda P, Kuete E, Dione N, Ngom II, Valles C, Bachar D, Raoult D, Lagier JC. 2019. Metagenomic and culturomic analysis of gut microbiota dysbiosis during Clostridium difficile infection. Scientific reports 9:1–8.

26. Sadowsky MJ, Staley C, Heiner C, Hall R, Kelly CR, Brandt L, Khoruts A. 2017. Analysis of gut microbiota–An ever changing landscape. Gut microbes 8:268–275.

27. Poirier D, Gervais P, Fuchs M, Roussy J-F, Paquet-Bolduc B, Trottier S, Longtin J, Loo VG, Longtin Y. 2019. Predictors of Clostridioides difficile Infection Among Asymptomatic, Colonized Patients: A Retrospective Cohort Study. Clin Infect Dis.

28. Terveer EM, Crobach MJ, Sanders IM, Vos MC, Verduin CM, Kuijper EJ. 2017. Detection of Clostridium difficile in feces of asymptomatic patients admitted to the hospital. J Clin Microbiol 55:403–411.

29. Song JH, Kim YS. 2019. Recurrent Clostridium difficile infection: risk factors, treatment, and prevention. Gut and liver 13:16.

30. Shin JH, Warren CA. 2019. Prevention and treatment of recurrent Clostridioides difficile infection. Curr Opin Infect Dis 32:482–489.

31. Seekatz AM, Wolfrum E, DeWald CM, Putler RK, Vendrov KC, Rao K, Young VB. 2018. Presence of multiple Clostridium difficile strains at primary infection is associated with development of recurrent disease. Anaerobe 53:74–81.

32. van Beurden YH, Nezami S, Mulder CJ, Vandenbroucke-Grauls C. 2018. Host factors are more important in predicting recurrent Clostridium difficile infection than ribotype and use of antibiotics. Clin Microbiol Infect 24:85. e1-85. e4.

33. Quraishi MN, Widlak M, Bhala Na, Moore D, Price M, Sharma N, Iqbal T. 2017. Systematic review with meta-analysis: the efficacy of faecal microbiota transplantation for the treatment of recurrent and refractory Clostridium difficile infection. Alimentary pharmacology & therapeutics 46:479–493.

34. Mullish BH, McDonald JA, Pechlivanis A, Allegretti JR, Kao D, Barker GF, Kapila D, Petrof EO, Joyce SA, Gahan CG. 2019. Microbial bile salt hydrolases mediate the efficacy of faecal microbiota transplant in the treatment of recurrent Clostridioides difficile infection. Gut 68:1791–1800.

35. Staley C, Kelly CR, Brandt LJ, Khoruts A, Sadowsky MJ. 2016. Complete microbiota engraftment is not essential for recovery from recurrent Clostridium difficile infection following fecal microbiota transplantation. MBio 7:e01965–16.

36. U.S. Food and Drug Administration. 2019. Important Safety Alert Regarding Use of Fecal Microbiota for Transplantation and Risk of Serious Adverse Reactions Due to Transmission of Multi-Drug Resistant Organisms.

37. Molinero N, Ruiz L, Sanchez B, Margolles A, Delgado S. 2019. Intestinal Bacteria interplay with bile and cholesterol metabolism: implications on host physiology. Frontiers in physiology 10:185.

38. Ridlon JM, Harris SC, Bhowmik S, Kang D-J, Hylemon PB. 2016. Consequences of bile salt biotransformations by intestinal bacteria. Gut microbes 7:22–39.

39. Ramírez-Pérez O, Cruz-Ramón V, Chinchilla-López P, Méndez-Sánchez N. 2018. The role of the gut microbiota in bile acid metabolism. Annals of hepatology 16:21–26.

40. Morrison DJ, Preston T. 2016. Formation of short chain fatty acids by the gut microbiota and their impact on human metabolism. Gut microbes 7:189–200.

41. Oliphant K, Allen-Vercoe E. 2019. Macronutrient metabolism by the human gut microbiome: major fermentation by-products and their impact on host health. Microbiome 7:91.

42. Schloissnig S, Arumugam M, Sunagawa S, Mitreva M, Tap J, Zhu A, Waller A, Mende DR, Kultima JR, Martin J. 2013. Genomic variation landscape of the human gut microbiome. Nature 493:45–50.

43. Dethlefsen L, Relman DA. 2011. Incomplete recovery and individualized responses of the human distal gut microbiota to repeated antibiotic perturbation. Proceedings of the National Academy of Sciences 108:4554–4561.

44. Lloyd-Price J, Mahurkar A, Rahnavard G, Crabtree J, Orvis J, Hall AB, Brady A, Creasy HH, McCracken C, Giglio MG. 2017. Strains, functions and dynamics in the expanded Human Microbiome Project. Nature 550:61.

45. De Filippis F, Vitaglione P, Cuomo R, Berni Canani R, Ercolini D. 2018. Dietary interventions to modulate the gut microbiome—how far away are we from precision medicine. Inflammatory bowel diseases 24:2142–2154.

46. Adalsteinsdottir SA, Magnusdottir OK, Halldorsson TI, Birgisdottir BE. 2018. Towards an individualized nutrition treatment: role of the gastrointestinal microbiome in the interplay between diet and obesity. Current obesity reports 7:289–293.

47. Inda ME, Broset E, Lu TK, de la Fuente-Nunez C. 2019. Emerging Frontiers in Microbiome Engineering. Trends Immunol.

48. Kumar M, Ji B, Zengler K, Nielsen J. 2019. Modelling approaches for studying the microbiome. Nature microbiology 4:1253–1267.

49. Zuñiga C, Zaramela L, Zengler K. 2017. Elucidation of complexity and prediction of interactions in microbial communities. Microbial biotechnology 10:1500–1522.

50. Bauer E, Thiele I. 2018. From network analysis to functional metabolic modeling of the human gut microbiota. MSystems 3:e00209–17.

51. Baldini F, Heinken A, Heirendt L, Magnusdottir S, Fleming RM, Thiele I. 2018. The Microbiome Modeling Toolbox: from microbial interactions to personalized microbial communities. Bioinformatics 35:2332–2334.

52. Hertel J, Harms AC, Heinken A, Baldini F, Thinnes CC, Glaab E, Vasco DA, Pietzner M, Stewart ID, Wareham NJ. 2019. Integrated Analyses of Microbiome and Longitudinal Metabolome Data Reveal Microbial-Host Interactions on Sulfur Metabolism in Parkinson’s Disease. Cell reports 29:1767-1777. e8.

53. Heinken A, Ravcheev DA, Baldini F, Heirendt L, Fleming RM, Thiele I. 2019. Systematic assessment of secondary bile acid metabolism in gut microbes reveals distinct metabolic capabilities in inflammatory bowel disease. Microbiome 7:75.

54. Seekatz AM, Aas J, Gessert CE, Rubin TA, Saman DM, Bakken JS, Young VB. 2014. Recovery of the gut microbiome following fecal microbiota transplantation. MBio 5:e00893–14.

55. Noronha A, Modamio J, Jarosz Y, Guerard E, Sompairac N, Preciat G, Daníelsdóttir AD, Krecke M, Merten D, Haraldsdóttir HS. 2018. The Virtual Metabolic Human database: integrating human and gut microbiome metabolism with nutrition and disease. Nucleic Acids Res 47:D614–D624.

56. Heirendt L, Arreckx S, Pfau T, Mendoza SN, Richelle A, Heinken A, Haraldsdottir HS, Wachowiak J, Keating SM, Vlasov V. 2019. Creation and analysis of biochemical constraint-based models using the COBRA Toolbox v. 3.0. Nature protocols 14:639.

57. Lovell D, Pawlowsky-Glahn V, Egozcue JJ, Marguerat S, Bähler J. 2015. Proportionality: a valid alternative to correlation for relative data. PLoS Comp Biol 11.

58. Davies DL, Bouldin DW. 1979. A cluster separation measure. IEEE transactions on pattern analysis and machine intelligence:224–227.

59. Allegretti JR, Kearney S, Li N, Bogart E, Bullock K, Gerber GK, Bry L, Clish CB, Alm E, Korzenik J. 2016. Recurrent Clostridium difficile infection associates with distinct bile acid and microbiome profiles. Alimentary pharmacology & therapeutics 43:1142–1153.

60. Abt MC, McKenney PT, Pamer EG. 2016. Clostridium difficile colitis: pathogenesis and host defence. Nature Reviews Microbiology 14:609–620.

61. Thanissery R, Winston JA, Theriot CM. 2017. Inhibition of spore germination, growth, and toxin activity of clinically relevant C. difficile strains by gut microbiota derived secondary bile acids. Anaerobe 45:86–100.

62. Lewis BB, Carter RA, Pamer EG. 2016. Bile acid sensitivity and in vivo virulence of clinical Clostridium difficile isolates. Anaerobe 41:32–36.

63. Crobach MJ, Vernon JJ, Loo VG, Kong LY, Péchiné S, Wilcox MH, Kuijper EJ. 2018. Understanding Clostridium difficile colonization. Clin Microbiol Rev 31:e00021–17.

64. Seekatz AM, Young VB. 2014. Clostridium difficile and the microbiota. The Journal of clinical investigation 124:4182–4189.

65. Hryckowian AJ, Pruss KM, Sonnenburg JL. 2017. The emerging metabolic view of Clostridium difficile pathogenesis. Curr Opin Microbiol 35:42–47.

66. Rodriguez C, Taminiau B, Korsak N, Avesani V, Van Broeck J, Brach P, Delmée M, Daube G. 2016. Longitudinal survey of Clostridium difficile presence and gut microbiota composition in a Belgian nursing home. BMC Microbiol 16:229.

67. Karlsson S, Lindberg A, Norin E, Burman LG, Åkerlund T. 2000. Toxins, butyric acid, and other short-chain fatty acids are coordinately expressed and down-regulated by cysteine in Clostridium difficile. Infect Immun 68:5881–5888.

68. Martin-Verstraete I, Peltier J, Dupuy B. 2016. The regulatory networks that control Clostridium difficile toxin synthesis. Toxins 8:153.

69. Blount KF, Shannon WD, Deych E, Jones C. Restoration of Bacterial Microbiome Composition and Diversity Among Treatment Responders in a Phase 2 Trial of RBX2660: An Investigational Microbiome Restoration Therapeutic, p ofz095. In (ed), Oxford University Press US,

70. Seekatz AM, Theriot CM, Rao K, Chang Y-M, Freeman AE, Kao JY, Young VB. 2018. Restoration of short chain fatty acid and bile acid metabolism following fecal microbiota transplantation in patients with recurrent Clostridium difficile infection. Anaerobe 53:64–73.

71. Weingarden AR, Chen C, Bobr A, Yao D, Lu Y, Nelson VM, Sadowsky MJ, Khoruts A. 2014. Microbiota transplantation restores normal fecal bile acid composition in recurrent Clostridium difficile infection. American Journal of Physiology-Gastrointestinal and Liver Physiology 306:G310–G319.

72. Khanna S, Montassier E, Schmidt B, Patel R, Knights D, Pardi DS, Kashyap PC. 2016. Gut microbiome predictors of treatment response and recurrence in primary Clostridium difficile infection. Alimentary pharmacology & therapeutics 44:715–727.

73. Staley C, Vaughn BP, Graiziger CT, Singroy S, Hamilton MJ, Yao D, Chen C, Khoruts A, Sadowsky MJ. 2017. Community dynamics drive punctuated engraftment of the fecal microbiome following transplantation using freeze-dried, encapsulated fecal microbiota. Gut microbes 8:276–288.

74. Gonzales M, Pepin J, Frost EH, Carrier JC, Sirard S, Fortier L-C, Valiquette L. 2010. Faecal pharmacokinetics of orally administered vancomycin in patients with suspected Clostridium difficile infection. BMC Infect Dis 10:363.

75. Schubert AM, Sinani H, Schloss PD. 2015. Antibiotic-induced alterations of the murine gut microbiota and subsequent effects on colonization resistance against Clostridium difficile. MBio 6:e00974–15.

76. Citron DM, Tyrrell KL, Merriam CV, Goldstein EJ. 2012. In vitro activities of CB-183,315, vancomycin, and metronidazole against 556 strains of Clostridium difficile, 445 other intestinal anaerobes, and 56 Enterobacteriaceae species. Antimicrobial agents and chemotherapy 56:1613–1615.

77. Jernberg C, Löfmark S, Edlund C, Jansson JK. 2010. Long-term impacts of antibiotic exposure on the human intestinal microbiota. Microbiology 156:3216–3223.

78. Tannock GW, Munro K, Taylor C, Lawley B, Young W, Byrne B, Emery J, Louie T. 2010. A new macrocyclic antibiotic, fidaxomicin (OPT-80), causes less alteration to the bowel microbiota of Clostridium difficile-infected patients than does vancomycin. Microbiology 156:3354–3359.

79. Louie TJ, Emery J, Krulicki W, Byrne B, Mah M. 2009. OPT-80 eliminates Clostridium difficile and is sparing of bacteroides species during treatment of C. difficile infection. Antimicrobial agents and chemotherapy 53:261–263.

80. Goldstein EJ, Babakhani F, Citron DM. 2012. Antimicrobial activities of fidaxomicin. Clin Infect Dis 55:S143–S148.

81. Vincent C, Manges AR. 2015. Antimicrobial use, human gut microbiota and Clostridium difficile colonization and infection. Antibiotics 4:230–253.

82. Sridharan GV, Choi K, Klemashevich C, Wu C, Prabakaran D, Pan LB, Steinmeyer S, Mueller C, Yousofshahi M, Alaniz RC. 2014. Prediction and quantification of bioactive microbiota metabolites in the mouse gut. Nature communications 5:1–13.

83. Pérez-Cobas AE, Artacho A, Ott SJ, Moya A, Gosalbes MJ, Latorre A. 2014. Structural and functional changes in the gut microbiota associated to Clostridium difficile infection. Frontiers in microbiology 5:335.

84. Riedel T, Wetzel D, Hofmann JD, Plorin SPEO, Dannheim H, Berges M, Zimmermann O, Bunk B, Schober I, Spröer C. 2017. High metabolic versatility of different toxigenic and non-toxigenic Clostridioides difficile isolates. Int J Med Microbiol 307:311–320.

85. Karlsson S, Burman LG, Åkerlund T. 2008. Induction of toxins in Clostridium difficile is associated with dramatic changes of its metabolism. Microbiology 154:3430–3436.

86. Khoruts A, Sadowsky MJ. 2016. Understanding the mechanisms of faecal microbiota transplantation. Nature reviews Gastroenterology & hepatology 13:508.

87. Brown JR-M, Flemer B, Joyce SA, Zulquernain A, Sheehan D, Shanahan F, O’Toole PW. 2018. Changes in microbiota composition, bile and fatty acid metabolism, in successful faecal microbiota transplantation for Clostridioides difficile infection. BMC gastroenterology 18:1–15.

88. Song Y, Garg S, Girotra M, Maddox C, von Rosenvinge EC, Dutta A, Dutta S, Fricke WF. 2013. Microbiota dynamics in patients treated with fecal microbiota transplantation for recurrent Clostridium difficile infection. PloS one 8.

89. Staley C, Kaiser T, Vaughn BP, Graiziger C, Hamilton MJ, Kabage AJ, Khoruts A, Sadowsky MJ. 2019. Durable Long-Term Bacterial Engraftment following Encapsulated Fecal Microbiota Transplantation To Treat Clostridium difficile Infection. mBio 10:e01586–19.

90. Jalanka J, Mattila E, Jouhten H, Hartman J, de Vos WM, Arkkila P, Satokari R. 2016. Long-term effects on luminal and mucosal microbiota and commonly acquired taxa in faecal microbiota transplantation for recurrent Clostridium difficile infection. BMC medicine 14:155.

91. Weingarden AR, Dosa PI, DeWinter E, Steer CJ, Shaughnessy MK, Johnson JR, Khoruts A, Sadowsky MJ. 2016. Changes in colonic bile acid composition following fecal microbiota transplantation are sufficient to control Clostridium difficile germination and growth. PloS one 11.

