## Supplemental Figures S1-S12 for "Computational Modeling of the Gut Microbiota Predicts Metabolic Mechanisms of Recurrent *Clostridioides difficile* Infection"

### Supplementary Information

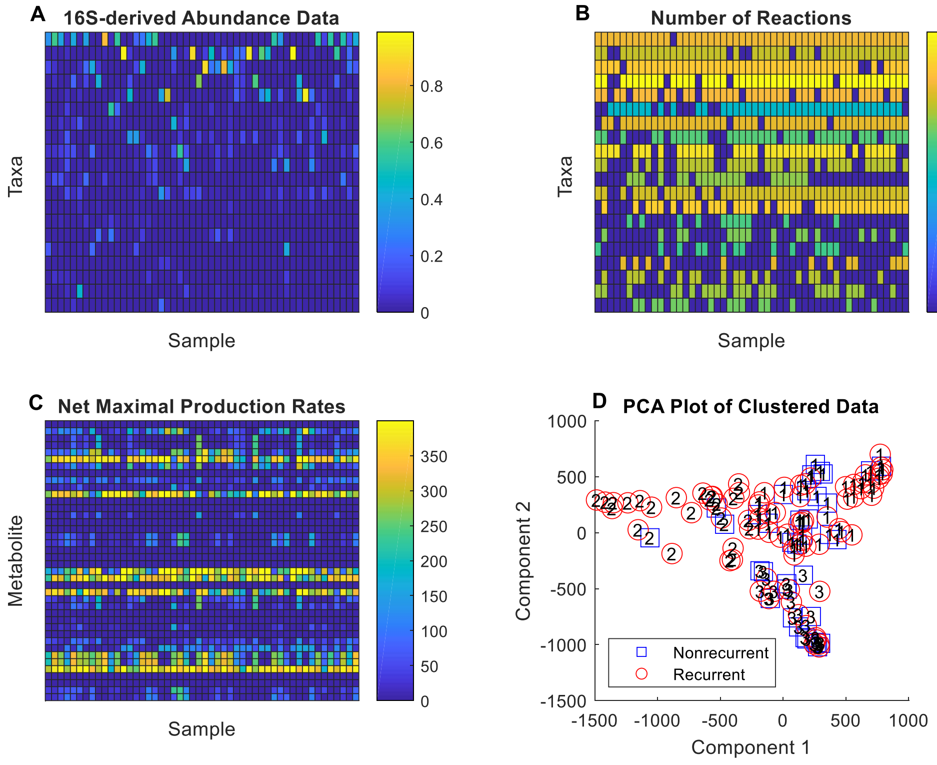


**Figure S1. Schematic representation of the community metabolic modeling work flow.** (A) Normalized taxa abundances were calculated from 16S rRNA data for CDI patient stool samples (PMC4847246). (B) Sample-specific community metabolic models were derived from the normalized taxa abundances using the Metagenomics Modeling Pipeline (mgPipe; PMC6596895) within the MATLAB Constraint-Based Reconstruction and Analysis (COBRA) Toolbox. (C) mgPipe computed the net maximal production rate (NMPC) of every exchanged metabolite for each sample model. (D) The NMPC simulation data was subjected to machine learning and statistical tests to extract information on CDI recurrence.

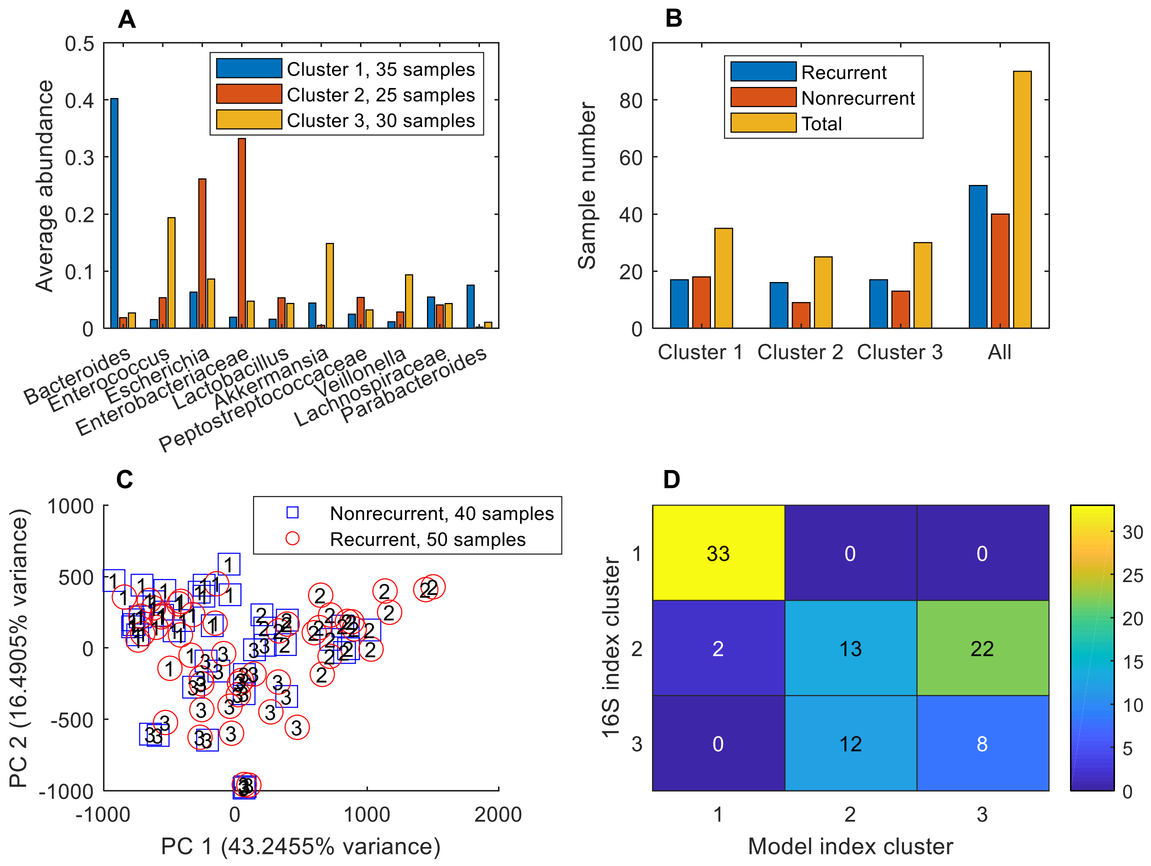


**Figure S2. Clustering of 90 index samples using model-processed abundance data.** (A) Average taxa abundances across the samples in each cluster for taxa which averaged at least 5% of the total abundance. (B) Number of recurrent, nonrecurrent and total samples in each cluster and all 90 index samples. None of the clusters contained a disproportionate number of recurrent samples (Fisher's exact test, $p > 0.25$). (C) PCA plot of the abundance data with each recurrent and nonrecurrent sample labeled by its associated cluster number. (D) Intersection between samples clustered based on model-processed abundance data and 16S-derived abundance data. The number in each box represents the number of shared samples between clusters.

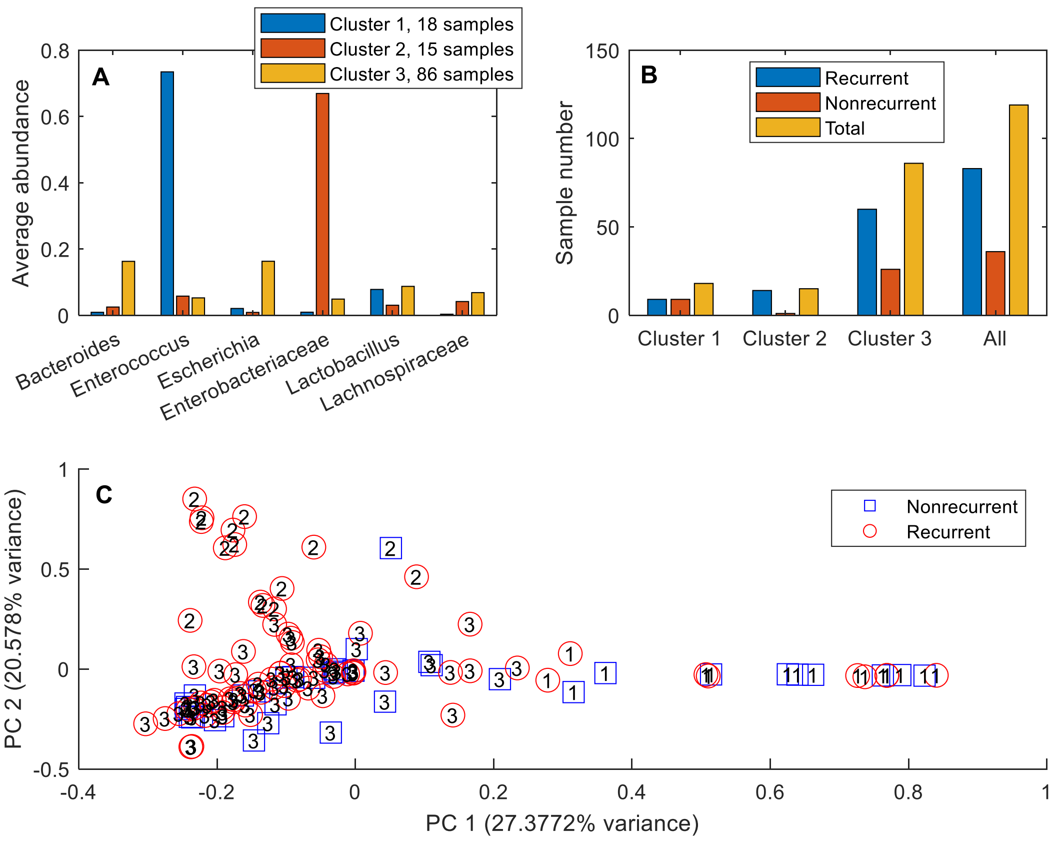


**Figure S3. Clustering of 119 post-index samples using 16S-derived abundance data.** (A) Average taxa abundances across the samples in each cluster for taxa which averaged at least 5% of the total abundance. (B) Number of recurrent, nonrecurrent and total samples in each cluster and all 119 post-index samples. Cluster 2 contained a disproportionate number of recurrent samples (14/15) compared to the cluster 1 (9/18; Fisher's exact test, $p = 0.009$). (C) PCA plot of the abundance data with each recurrent and nonrecurrent sample labeled by its associated cluster number.

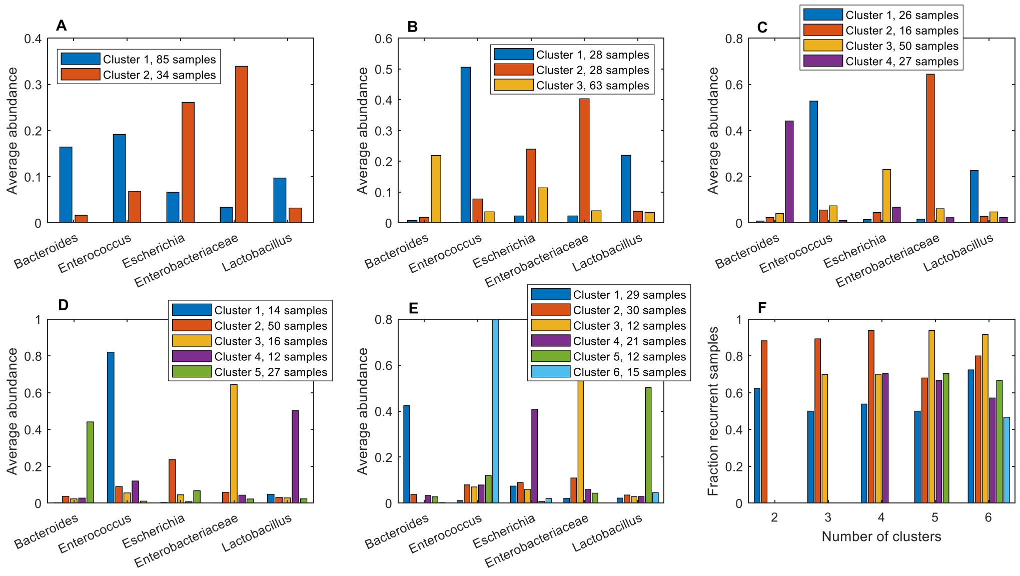


**Figure S4. Clustering of 119 post-index samples with 2 to 6 total clusters using model-processed abundance data.** Average taxa abundances across the samples in each cluster for the 5 most abundant taxa across all samples: (A) 2 clusters; (B) 3 clusters; (C) 4 clusters; (D) 5 clusters; and (E) 6 clusters. (F) Fraction of recurrent samples in each cluster for 2 to 6 total clusters. The following clusters had a disproportionate number of recurrent samples based on Fisher's exact test: 2 total clusters, cluster 1 versus cluster 2 ($p = 0.007$); 3 total clusters, cluster 2 versus cluster 1 ($p = 0.003$); 4 total clusters, cluster 2 versus cluster 1 ($p = 0.007$); 5 total clusters, cluster 3 versus cluster 1 ($p = 0.012$); 6 total clusters, cluster 3 versus cluster 6 ($p = 0.019$), cluster 2 versus cluster 6 ($p = 0.039$).

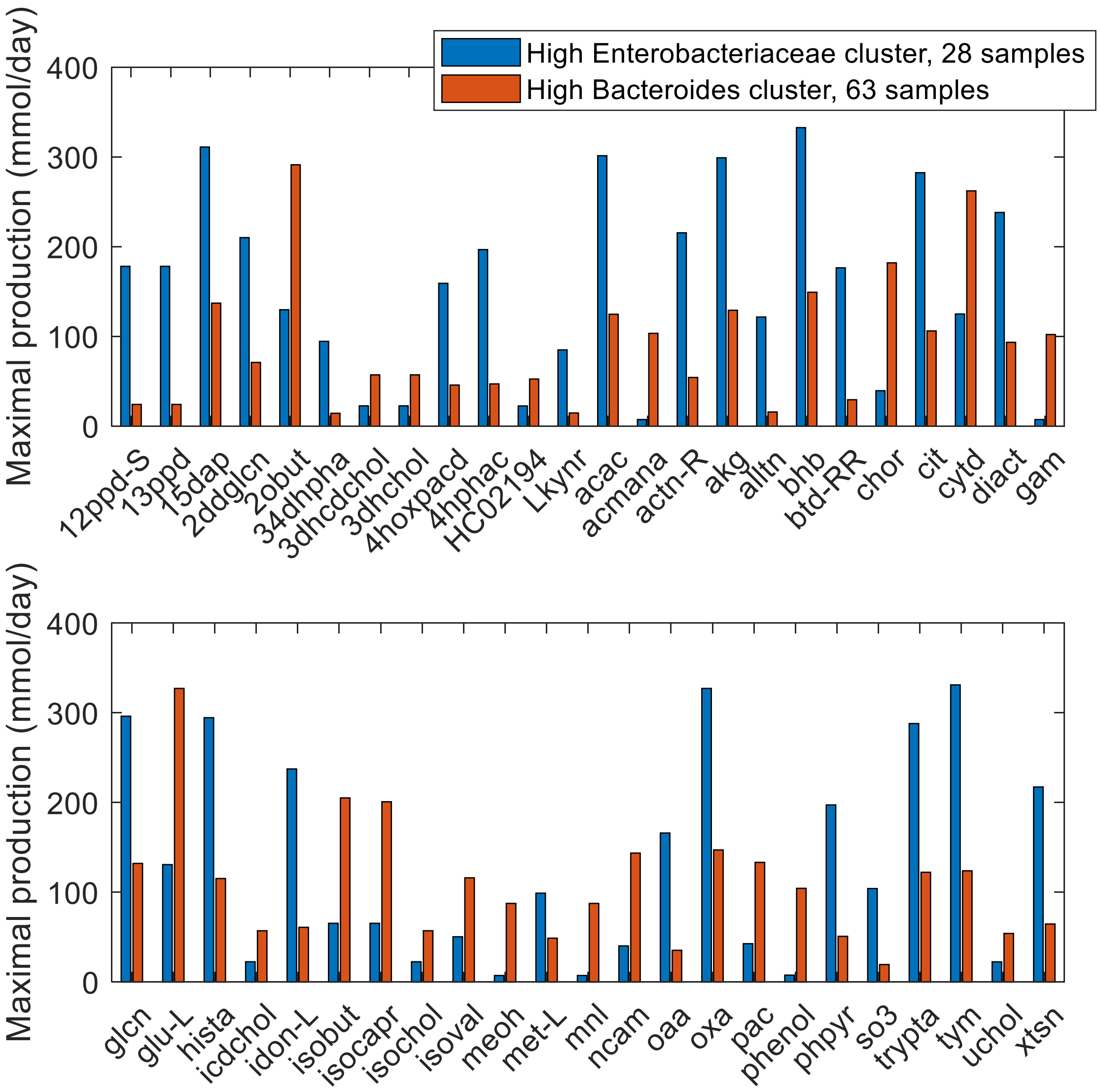


**Figure S5. Differentially produced metabolites in the high *Enterobacteriaceae* and high *Bacteroides* clusters generated from 119 model-processed post-index samples.** Significant differences in metabolite production rates were determined by applying the Wilcoxon rank sum test ($p < 0.05$) to each metabolite across all samples in the two clusters. In addition to being statistically different, each metabolites shown had an average production rate > 50 mmol/day in at least one cluster and average production rates that differed between the clusters by at least 100%. Metabolite abbreviations are taken from the VMH database (www.vmh.life). Full metabolite names, their associated metabolic pathways and numeric values for their average production rates in each cluster are given in Table S6.

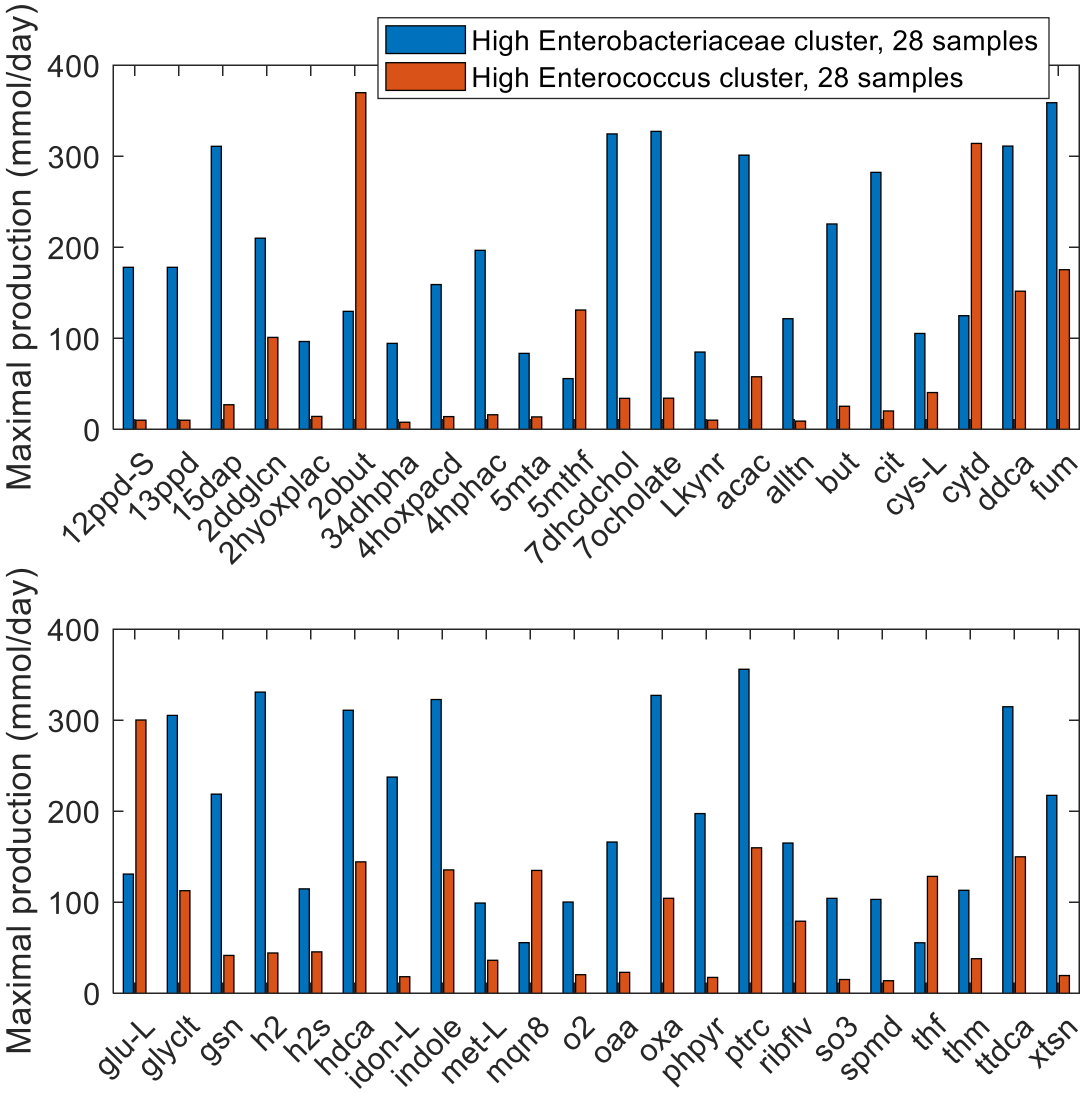


**Figure S6. Differentially produced metabolites in the high *Enterobacteriaceae* and high *Enterococcus* clusters generated from 119 model-processed post-index samples.** Significant differences in metabolite production rates were determined by applying the Wilcoxon rank sum test ($p < 0.05$) to each metabolite across all samples in the two clusters. In addition to being statistically different, each metabolites shown had an average production rate > 50 mmol/day in at least one cluster and average production rates that differed between the clusters by at least 100%. Metabolite abbreviations are taken from the VMH database (www.vmh.life). Full metabolite names, their associated metabolic pathways and numeric values for their average production rates in each cluster are given in Table S6.

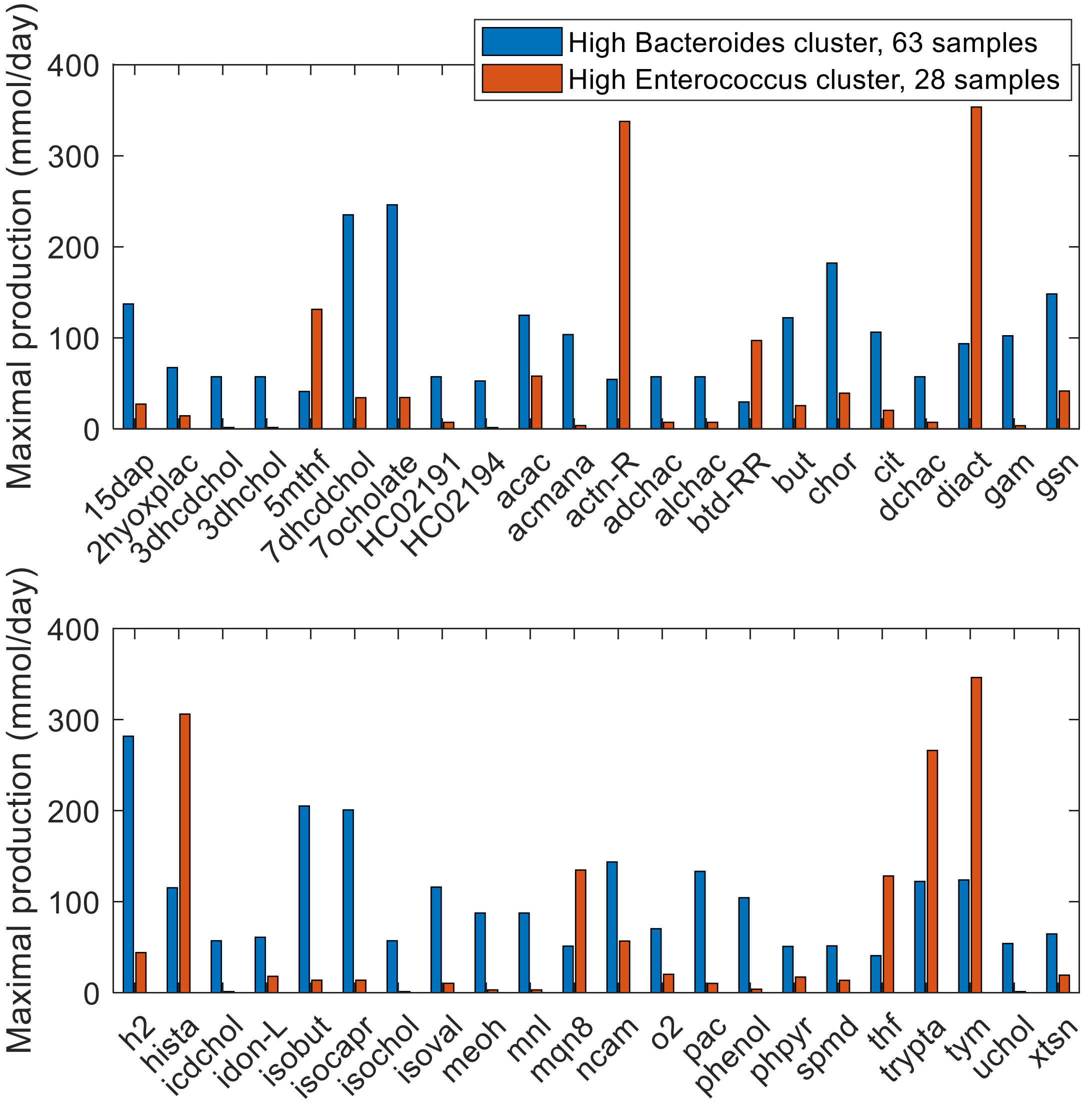


**Figure S7. Differentially produced metabolites in the high *Bacteroides* and high *Enterococcus* clusters generated from 119 model-processed post-index samples.** Significant differences in metabolite production rates were determined by applying the Wilcoxon rank sum test ($p < 0.05$)

to each metabolite across all samples in the two clusters. In addition to being statistically different, each metabolites shown had an average production rate > 50 mmol/day in at least one cluster and average production rates that differed between the clusters by at least 100%. Metabolites abbreviations are taken from the VMH database (www.vmh.life). Full metabolite names, their associated metabolic pathways and numeric values for their average production rates in each cluster are given in Table S6.

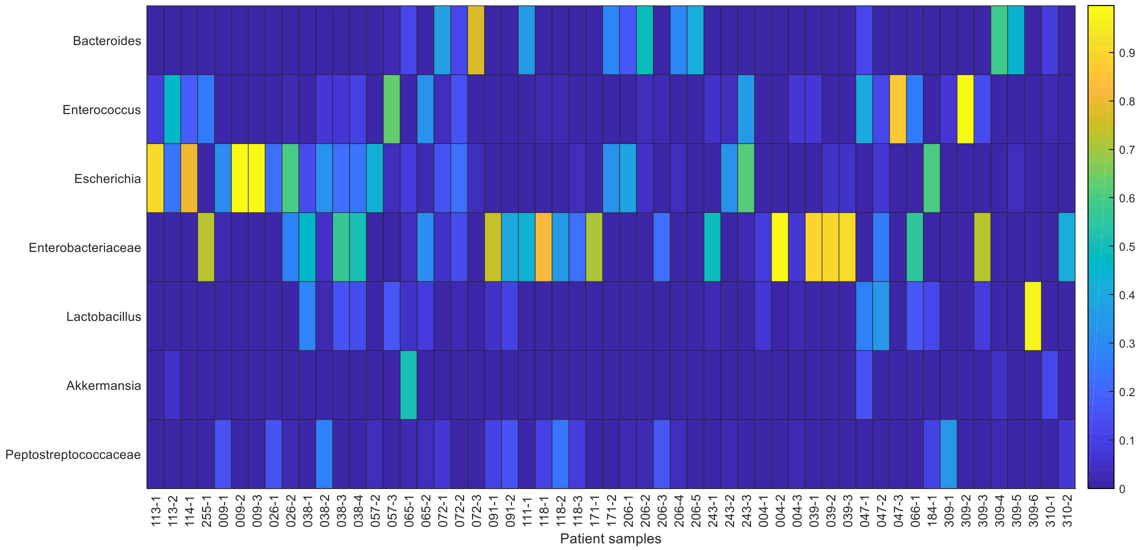


**Figure S8. Abundances of the top 7 taxa for all post-index samples of the 22 patients in the high *Enterobacteriaceae* (HEb) group.** All post-index samples from the 22 patients with at least one sample in the high *Enterobacteriaceae* (HEb) cluster were grouped to generate the HEb group of 55 samples. The samples are denoted as XXX-Y where XXX is the patient ID and Y is the post-index sample number of that patient.

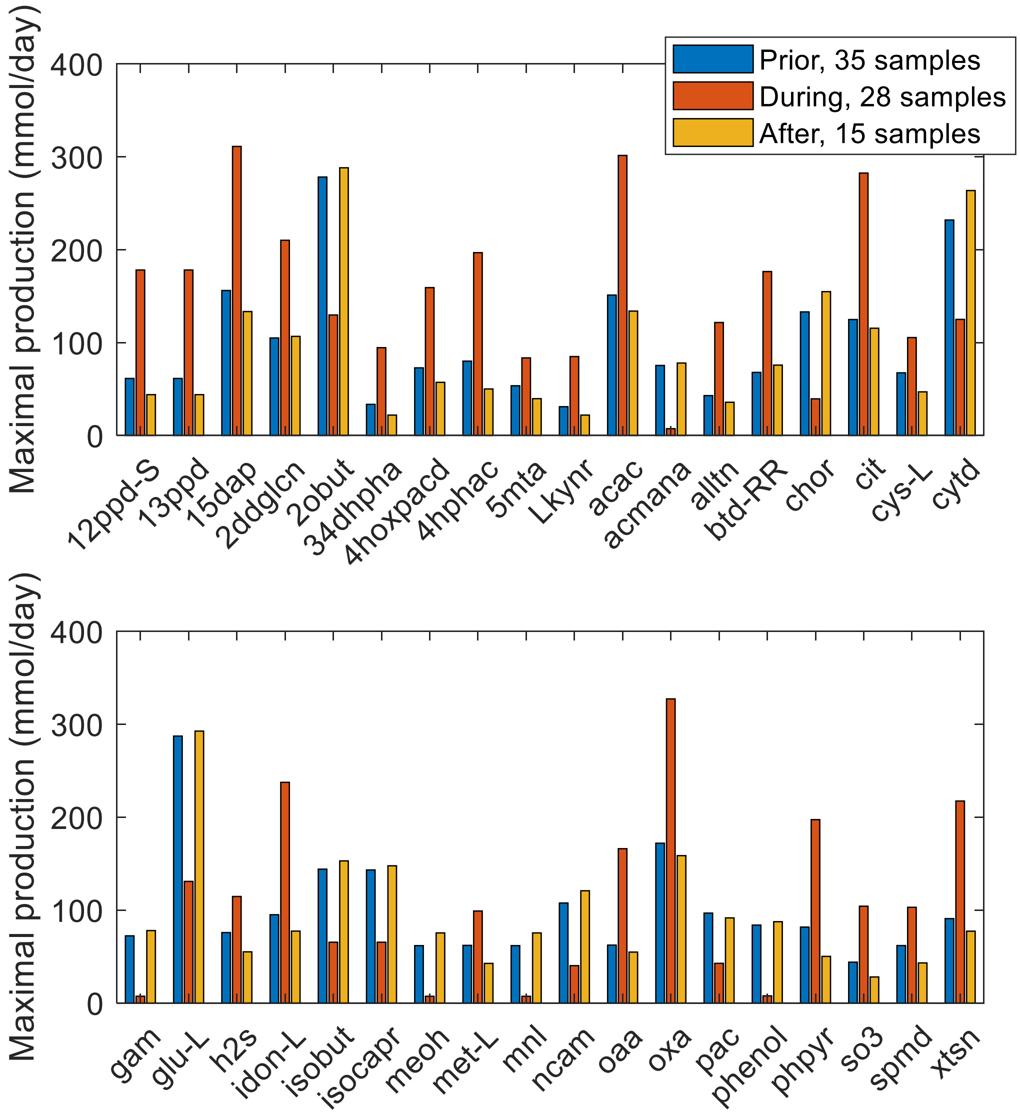


**Figure S9. Differentially produced metabolites of the 22 patients in the expanded high *Enterobacteriaceae* (HEb) group.** The HEb group was expanded to contain pre-index and index samples to generate a dataset of 78 samples. These samples were partitioned into 35 samples prior to patients entering the HEb cluster, 28 samples during patient presence in the HEb cluster and 15 samples after patients left the HEb cluster. Significant differences in metabolite production rates were determined by applying the Wilcoxon rank sum test ($p < 0.05$) to each metabolite across all samples in the two groups compared (e.g. pre-HEb vs. HEb, pre-HEb vs. post-HEb, pre-HEb vs.

post-HEb). In addition to being statistically different between at least two sample groups, each metabolites shown had an average production rate > 50 mmol/day in at least one group and average production rates that differed between the two groups by at least 100%. Metabolites abbreviations are taken from the VMH database ([www.vmh.life](www.vmh.life)). Full metabolite names and their associated metabolic pathways are given in Table S6.

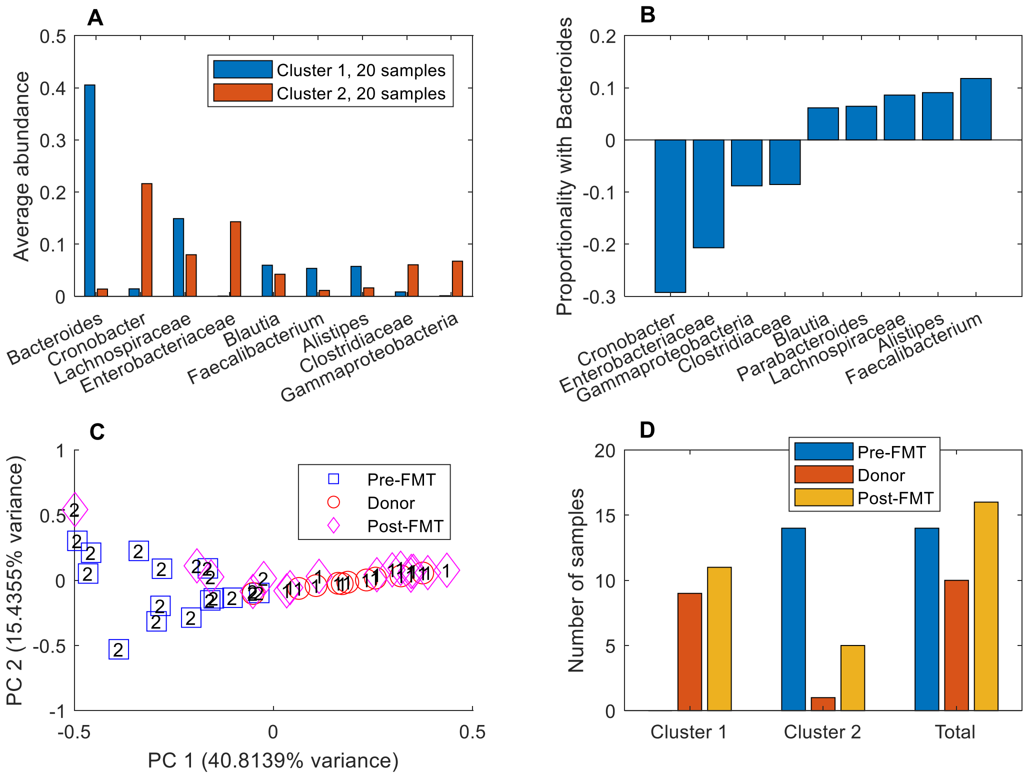


**Figure S10. Clustering of 40 FMT samples using 16S-derived abundance data.** (A) Average taxa abundances across the samples in each cluster for taxa which averaged at least 5% in at least one cluster. (B) Correlations between *Bacteroides* and other taxa calculated from all samples as measured by the proportionality coefficient ρ. The 9 taxa with the largest |ρ| values are shown. (C) PCA plot of the abundance data with each pre-FMT, donor and post-FMT sample labeled by its associated cluster number. (D) Number of pre-FMT, donor and post-FMT samples in each cluster and all 40 FMT samples. Cluster 2 contained a disproportionately large number of pre-FMT samples (14/20) compared to the cluster 1 (0/20; p < 0.00001) and the entire FMT dataset (14/40; p = 0.0014).

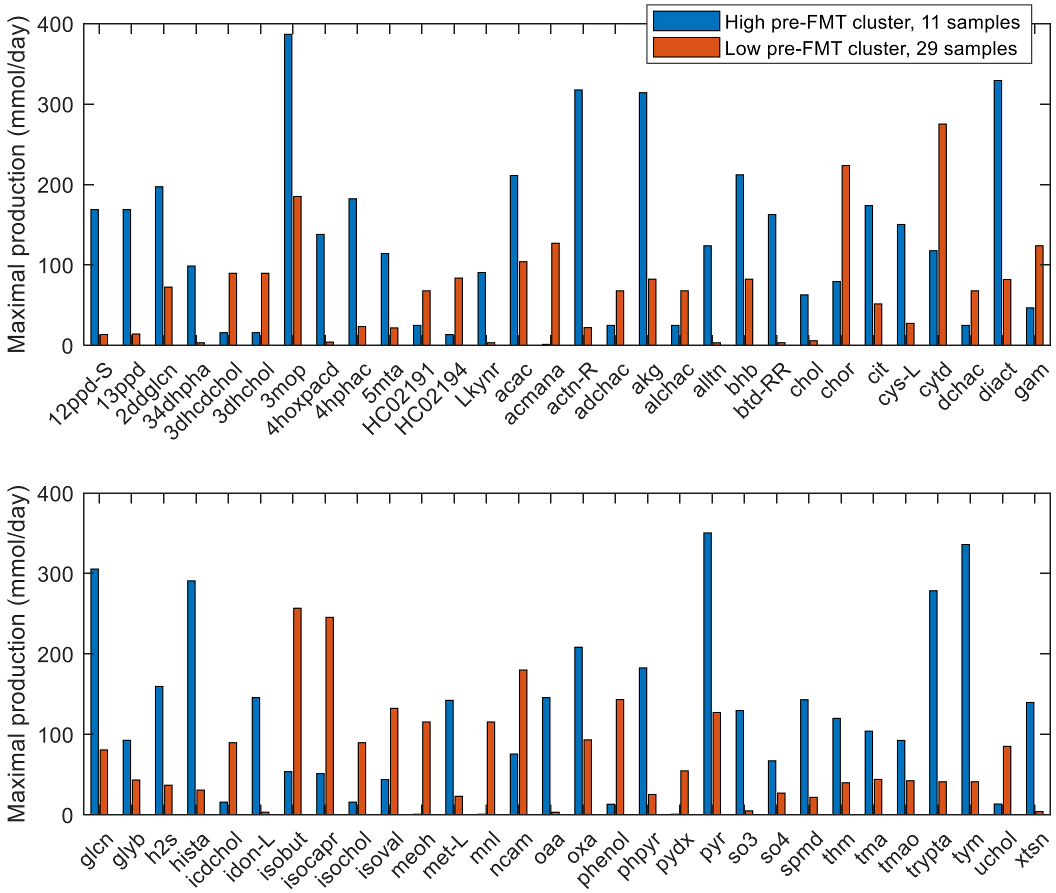


**Figure S11. Differentially produced metabolites in the high *Enterobacteriaceae* and high *Bacteroides* clusters generated from 40 model-processed FMT samples.** Significant differences in metabolite production rates were determined by applying the Wilcoxon rank sum test ($p < 0.05$) to each metabolite across all samples in the two clusters. In addition to being statistically different, each metabolites shown had an average production rate > 50 mmol/day in at least one cluster and average production rates that differed between the clusters by at least 100%. Metabolites abbreviations are taken from the VMH database (www.vmh.life). Full metabolite names, their associated metabolic pathways and numeric values for their average production rates in each cluster are given in Table S7.

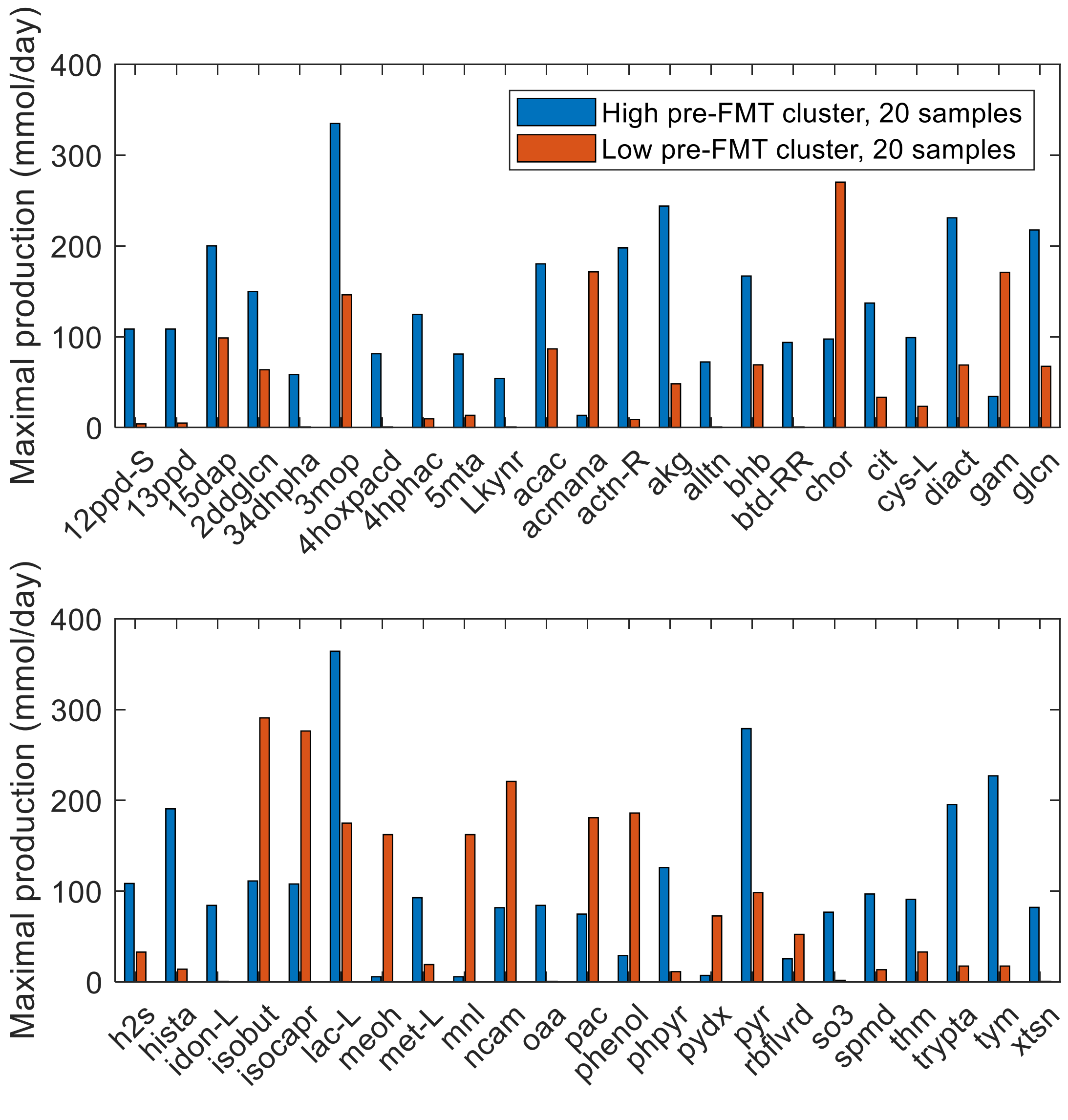


**Figure S12. Differentially produced metabolites in the high *Enterobacteriaceae* and high *Bacteroides* clusters generated from 40 FMT samples.** Significant differences in metabolite production rates were determined by applying the Wilcoxon rank sum test ($p < 0.05$) to each metabolite across all samples in the two clusters. In addition to being statistically different, each metabolites shown had an average production rate > 50 mmol/day in at least one cluster and average production rates that differed between the clusters by at least 100%. Metabolites abbreviations are taken from the VMH database (www.vmh.life). Full metabolite names and their associated metabolic pathways are given in Table S6 and S7.
